# Cyanophage CP12 Rewires Host Carbon Regulation through Interface Remodeling and Redox Buffering

**DOI:** 10.64898/2026.09.10.750766

**Authors:** Scott Widmann, Ruonan Wu, Malio Nelson, Daniel Mejia-Rodriguez, Suman Samantray, Reece Neff, Song Feng, Xiaolu Li, Jordan Rozum, August George, Paul Rigor, Pavlo Bohutskyi, Noelani Boise, Hoshin Kim, Lindsey N. Anderson, Wei-Jun Qian, Esther Olaleye, Kehinde Idowu, Amity Andersen, Margaret S. Cheung

## Abstract

Picocyanobacteria drive ocean carbon fixation, and cyanophages reshape host metabolism during infection. In cyanobacteria, the intrinsically disorder Calvin cycle protein 12 (CP12) assembles glyceraldehyde-3-phosphate dehydrogenase (GAP2) and phosphoribulokinase (PRK) into the inhibitory dark complex, yet how phage CP12 homologs modulate redox-sensitive partners remains unclear. Here, we examined viral CP12 across sequence, structure, and post-translational modification (PTM) complexities to resolve the mechanistic remodeling of dark-complex assembly and regulation. Protein-family analysis of CP12 homologs across diverse lineages showed that viral CP12 preserves interface-dominated positions while shifting partner-facing chemistry toward charged and geometry-modulating features. Matched molecular dynamics simulations (MD) of *Prochlorococcus* MED4 and cyanophage P-HM2 showed that phage CP12 preserves assembly while strengthening PRK-facing contacts and reducing GAP2-facing interface burial. MED4 redox proteomics identified coordinated cysteine oxidation across CP12 and GAP2 under light disturbance, guiding MD to explore thiol-PTM states. Conformational divergence increased with PTM load and localized mainly to GAP2 modifications. Thiol PTM at GAP2 imposed the largest CP12 binding-energy cost, which P-HM2 CP12 buffered, yielding smaller comparative binding-energy penalties than host CP12. These findings link sequence-driven chemistry to interface dynamics and redox PTM responsiveness to light, defining phage CP12 as a regulatory mimetic that may retune host carbon regulation during infection.

## Introduction

Viruses can reshape microbial metabolism by carrying auxiliary viral genes (AVGs) that are expressed during infection, altering host physiology and resource allocation with consequences for ecosystem-relevant processes^1–3^. AVGs are usually discussed in relation to auxiliary metabolic genes (AMGs), which encode enzymes that supplement or reroute host reactions in photosynthesis, carbon fixation, and nutrient acquisition^4–7^. Their discovery broadened understanding of virus-host interactions beyond lysis, yet metabolic rerouting is only one way viral proteins alter the host state. Other AVGs, known as auxiliary regulatory genes (AReGs), instead encode regulatory proteins thought to change how host protein complexes assemble, respond to environmental cues, and regulate enzyme activity^6,8^. Unlike viral metabolic enzymes, AReG-encoded proteins must remain compatible with host partners to form functional complexes while altering their behavior. Substitutions relative to host homologs may therefore reshape the stability, conformation, or partner engagement of host assemblies, but these effects are difficult to infer from sequence annotation alone.

Calvin cycle protein 12 (CP12) is a useful case for studying such remodeling. The light-dependent Calvin-Benson cycle fixes carbon in cyanobacteria, algae, and plants, and must be coordinated with electron transport in the photosynthetic process and cellular redox state^1,2^. A recent light disturbance study further shows that cyanobacterial carbon metabolism is regulated through rapid redox-proteomic responses that are separated from slower transcriptional changes^9^. CP12 is a redox-regulated intrinsically disordered protein that assembles with phosphoribulokinase (PRK) and glyceraldehyde-3-phosphate dehydrogenase (GAP2) into the inhibitory dark complex^3,4^. During light-to-dark perturbations, it promotes this assembly, suppressing carbon fixation as photosynthetic reducing power declines^5^. With an estimated global population of around 3 × 10^27^ cells, *Prochlorococcus* is the most abundant photosynthetic organism on Earth and, with *Synechococcus*, contributes roughly 25% of ocean net primary production^6^, linking Calvin-Benson regulation to global carbon cycling. The alpha-cyanobacterium *Prochlorococcus marinus* MED4 is a minimal photoautotroph and a tractable model for photosynthetic carbon regulation, with a dark complex of two GAP2 homotetramers and two PRK homodimers bridged by four CP12 molecules^1,7^. CP12 homologs also occur in cyanophages, including the T4-like P-HM2 infecting MED4. Our recent genome-scale metabolic model predicted that phage CP12 mediated Calvin-Benson inhibition and redirected flux from carbon fixation toward the pentose phosphate pathway and nucleotide metabolism, with this system-level impact validated by nitrogen-dependent growth in the P-HM2 CP12-supplemented cyanobacterial culture^10^. Because cyanophages lyse an estimated 20% of the cyanobacterial population daily^8^ and carry AMGs that reshape host photosynthesis and central carbon metabolism^11^, phage CP12 homologs could influence host carbon regulation at ecologically meaningful scales. These homologs are substantially divergent from host CP12 sequences^11–13^, and it remains unknown whether phage CP12 simply mimics host CP12, weakens the host dark complex, or preserves the complex while changing its regulatory interface dynamics.

Annotation can identify a viral protein as CP12-like, but not whether it preserves host-like binding, rewires partner-specific contacts, or changes redox-dependent regulation. Static structure prediction, including AlphaFold^14^, provides valuable hypotheses, but CP12 is intrinsically flexible, undergoes disorder-to-order transitions upon binding, and acts through dynamic interactions with multiple partners, rendering static prediction incomplete^1,5^. Molecular dynamics (MD) simulations resolve how substitutions affect conformational sampling, contact persistence, interface burial, solvent exposure, and energetic redistribution over time^15,16^. Atomistic simulations of large assemblies remain computationally expensive, however, and cannot fully capture the natural diversity of CP12 or all cellular conditions^17,18^, highlighting the need for integrative approaches linking evolutionary variation with mechanistic insight.

Post-translational modifications (PTMs) add a rapid regulatory layer, altering enzyme activity and assembly without new transcription or translation. In photosynthetic metabolism, cysteine redox PTMs are a major mechanism coupling enzyme function to light-dependent redox conditions^19,20^. Dark complex regulation is discussed largely in terms of CP12 cysteine oxidation and reduction, with oxidized CP12 promoting assembly and reduced CP12 favoring dissociation^5^. However, MED4 CP12 lacks the canonical N-terminal cysteine pair found in many four-cysteine CP12 proteins, consistent with other reported cyanobacterial homologs^21^, suggesting a canonical two-disulfide switching in CP12 alone may not explain how it conditionally orders and regulates assembly. GAP2 and PRK are themselves redox-sensitive, and modifications beyond CP12 disulfide formation may contribute to dark complex behavior^22^. It remains unclear whether regulation is driven primarily by CP12 cysteines or by coordinated redox states across CP12 and its partners, and whether viral CP12 alters the complex’s response to redox perturbation.

Here, we address these gaps with a cross-scale framework connecting protein-family-wide sequence features to model-system structural dynamics and redox proteomics. To capture the sequence diversification accumulated over evolutionary timescales of host-phage adaptation, we integrated Salish Sea metagenome^23^ -recovered and reference database-retrieved CP12 homologs across microbial lineages. These sequences were first assessed for protein-family-wide physicochemical signatures at interface-linked positions. We then tested whether these signatures translate into fast regulatory timescales in metabolic processes through altered contact dynamics, interface burial, and energetic redistribution in paired MED4 and P-HM2 simulations. In parallel, we conducted the experiments of MED4 redox proteomics under light variation at both dawn and dusk, then identified responsive cysteine oxidation states. The redox state of cysteine residues, as reflected by PTM levels, can change bidirectionally between light and dark phases of the day, independent of protein abundance. We used the redox proteomics results to seed high-throughput combinatorial PTM simulations testing whether the same residues differentially affect host and phage CP12-containing complexes. This framework reveals phage CP12 as an AReG product that preserves host dark complex assembly while retuning its physicochemical and redox-sensitive dynamics, providing a cross-scale approach for identifying viral proteins that remodel host metabolism through protein-complex regulation with PTM, expanding understanding of viral metabolic control beyond lysis and direct host metabolic flux rerouting.

## Results

### CP12 protein family diversity reveals interface-structured chemical constraints

To connect sequence diversity to mechanistic interpretation across the CP12 protein family, we compiled dereplicated metagenome-derived (n = 123) and database-deposited (n = 1,298) CP12 homologs spanning bacterial, archaeal, eukaryotic, and viral lineages. Metagenome-derived CP12 broadened the observed ranges of cysteine number, cysteine spacing, and acidic residue fraction, three features relevant to redox sensitivity and charge patterning (**Fig. S1a-c**, **Supplementary Results**). This was not explained by incomplete sequence recovery, as full-length and partial metagenome-derived sequences showed overlapping distributions across the same properties (**Fig. S1a-c**). Metagenome-derived CP12 also expanded phylogenetic breadth, adding viral and non-cyanobacterial diversity to the public database that is currently over-represented by cyanobacterial homologs (**Fig. S1d**).

We projected all 1,421 sequences into the CP12 Hidden Markov Model (HMM: PF02672) for residue-level comparison. The median occupancy of each HMM position was 0.964 (**Fig.1b**). We linked HMM-aligned variation to reference-guided interface classes using two paired model systems, cyanobacterial host *Prochlorococcus* MED4 and its infecting cyanophage P-HM2, which serve as representative host-phage CP12 counterparts throughout this study. Both mapped extensively to the HMM, with MED4 CP12 aligning 91.9% of residues (97.1% column occupancy) and P-HM2 CP12 aligning 95.7% (95.7% occupancy) (**Fig.1a**). Each HMM position was assigned as a system-shared interface, a system-specific interface, or non-interface based on whether MED4 and P-HM2 CP12 residues contacted PRK and/or GAP2 in the oxidized dark complex (**Fig.1b**). We found 67 of 70 HMM positions were interface classes: 50 system-shared, 9 MED4-specific, 8 P-HM2-specific, and 3 non-interface, showing CP12 is interface-dominated despite lineage diversity. Across the 1,421 sequences, strong cysteine enrichment was confined to four positions: 57 and 66 (fractions 0.98 and 0.95) and 16 and 26 (0.71 each) with cysteine fraction at other positions below 0.005 (**Fig. 1b**). Positions 57 and 66 were system-shared GAP2-facing interfaces; 16 and 26 were MED4-specific and system-shared interfaces, both PRK-contacting.

**Figure 1.**
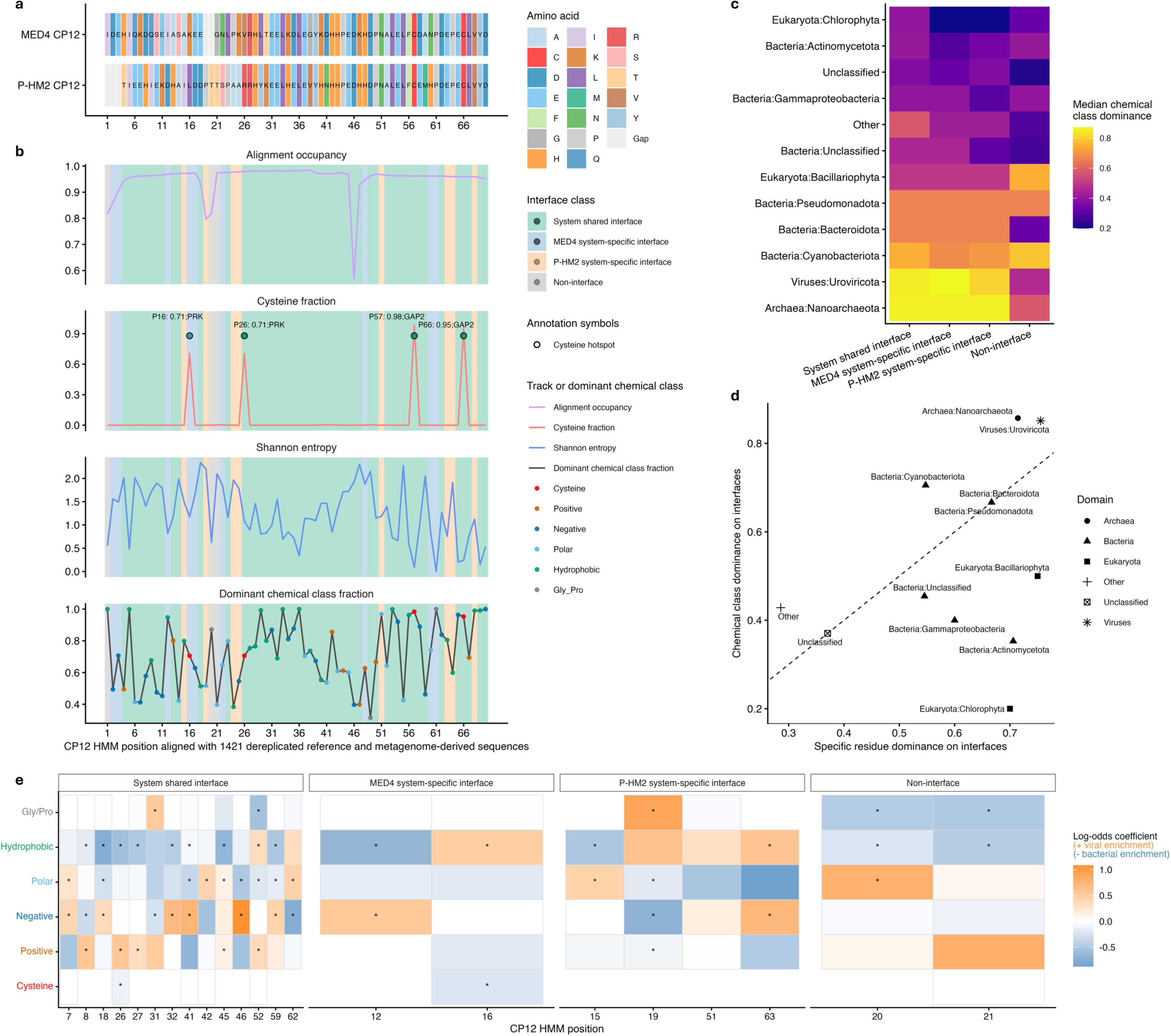
HMM-aligned CP12 conservation profiles. **a** Alignment of the two reference sequences (P-HM2 CP12 and MED4 CP12) across the aligned CP12 HMM positions. Tiles are colored by amino-acid identity (20-residue palette), with gaps shown in light gray; letters indicate aligned residues at occupied positions. **b** Protein family-wide per-position conservation tracks across 1,421 dereplicated CP12 sequences. Background shading marks interface context inferred from the two model systems (shared interface, MED4-specific interface, HM2-specific interface, non-interface). Dominant chemical-class points are overlaid on the dominance track. Annotated cysteine highlighted with contacting partners and its cysteine fraction. **c** Median chemical-class dominance by lineage and interface category. **d** Lineage summary scatter comparing median residue-identity dominance (x-axis) versus median chemical-class dominance (y-axis) across interface positions; the dashed diagonal indicates equality (chemistry-level conservation equals residue-level conservation). Points above the diagonal indicate stronger conservation at the chemical-class level than at exact residue identity. **e** Logistic-regression coefficients mapped to CP12 HMM positions and chemical classes. Positive coefficients (red) indicate viral enrichment, whereas negative coefficients (blue) indicate bacterial enrichment. Columns are HMM positions grouped by reference-guided interface class. Displayed positions passed position-level Fisher’s exact test with FDR correction (FDR < 0.01) and were ranked by maximum absolute coefficient. Asterisks highlighted chemical class-specific differences significant at the feature level (FDR < 0.01), based on Fisher’s exact tests comparing the presence versus absence of each chemical class between viral and bacterial sequences.

The alignment also resolved where CP12 is constrained at residue identity and chemical class levels. Shannon entropy was broadly distributed (**Fig. 1b**), exceeding 1 at 57 of 70 columns, 2 at 32, and 3 at only 6, indicating widespread moderate variation. The C-terminal region, enriched in GAP2-interface positions, showed lower entropy than the PRK-contacting N-terminus. Chemical class conservation was often strong. At 56 of 70 positions, a single class accounted for ≥50% of residues (mean dominant-class fraction 0.71) (**Fig. 1b**). We then tested the strength of the observed enrichment patterns using a null model randomly reassigning amino acids to chemical classes (**Supplementary Methods and Results**) and also found that interface classes may follow distinct constraint patterns. Residues at the P-HM2-specific HMM positions were significantly enriched for both dominant class preference and within-class variation, while residues at the HMM annotated as MED4-specific and shared interfaces showed significant substitution tolerance but weaker class preference (**Fig. S2**). Taxon-resolved analyses further showed that *Uroviricota* and Nanoarchaeota showed stronger chemical class dominance across interface classes (**Fig. 1c**) and stronger class preservation than bacterial and eukaryotic lineages (**Fig. 1d**). CP12 variation therefore remains structured by interface class across broad taxonomic diversity.

### Sequence-encoded chemical features distinguish viral and bacterial CP12

We next examined whether chemical-class patterns distinguish viral from bacterial CP12. Adjusted pairwise alignment of MED4 and P-HM2 CP12 using HMM positions revealed 64 gap-free aligned positions with 28 substitutions. Substitutions altered volume the most (net +9.43Å^3^ from MED4 to P-HM2), whereas hydrophobicity (Kyte-Doolittle) varied narrowly (mean +0.14) and charge changes were sparse (mean - 0.11) (**Fig. S3**). The broader protein-family analysis revealed a strong position-structured chemical-class contrast between viral and bacterial CP12s. Using HMM-position-resolved chemical-class features, a logistic regression classifier separated viral and bacterial sequences under cross-validation, achieving a mean Receiver Operating Characteristic Area Under the Curve (ROC AUC) of 0.995 and PR AUC of 0.850 (**Fig. S4a** and **4b**), with precision limited mainly by class imbalance. At a 0.5 threshold, it correctly identified 47 of 49 viral sequences (95.9% recall) and 1,299 of 1,316 bacterial sequences (**Fig. S4c**). Sequence-encoded chemical features therefore capture viral-like CP12 signatures, which we use here for ranking rather than strict binary classification.

To identify which classes at which positions drive this separation, we combined logistic-regression coefficients with Fisher’s exact tests and false discovery rate (FDR) correction. Bacterial CP12 were often characterized by hydrophobic or polar features at interface-linked positions, including shared positions 8, 18, 26, 27, 32, 41, 45, 46, 52 and 59, MED4-specific position 12, and P-HM2-specific positions 15 and 19 (**Fig. 1e**). Viral CP12 more frequently favored charged classes, with a few gains in polar or Gly/Pro classes (positive charge at shared positions 8, 26, 27, 45, 52; negative charge at shared positions 18, 32, 41, 46, 59, and MED4-specific position 12), suggesting viral CP12 may tune local contact geometry at selected sites. Cysteine contributed little to the classifier signal except at positions 16 and 26, consistent with cross-lineage conservation. These cysteines are absent in MED4 CP12 but reflect broader bacterial features, so patterns should be read as classifier-weighted signatures, not one-to-one substitutions. These signatures suggest viral CP12 remodels interface-linked physicochemical states, shifting from bacterial hydrophobic or polar features toward charged or geometry-modulating features. Because such mechanisms cannot be resolved from sequence alone, we next used paired MED4 and P-HM2 dark complex molecular dynamics (MD) simulations.

### P-HM2 CP12 preserves the dark complex binding arrangement across fully reduced and reference oxidized states

We built matched atomistic homology models of the MED4 and P-HM2 dark complexes using the *Thermosynechococcus vestitus* GAPDH-CP12-PRK cryo-EM structure as a template (**Fig. 2a-b**). Both models shared identical MED4 GAP2 and PRK subunits, differing only in CP12 identity (**Supplementary Methods; Tables S1 and S2**). Quality assessment supported stereochemical validity (**Fig. S5a**), and backbone root mean square deviation (RMSD) reached stable plateaus across three replicate 100 ns simulations (**Fig. S5b**). Thermodynamic integration indicated CP12 disulfide formation was energetically favorable in both complexes (**Fig. S5c; Supplementary Methods and Results**), supporting use of these models.

**Figure 2.**
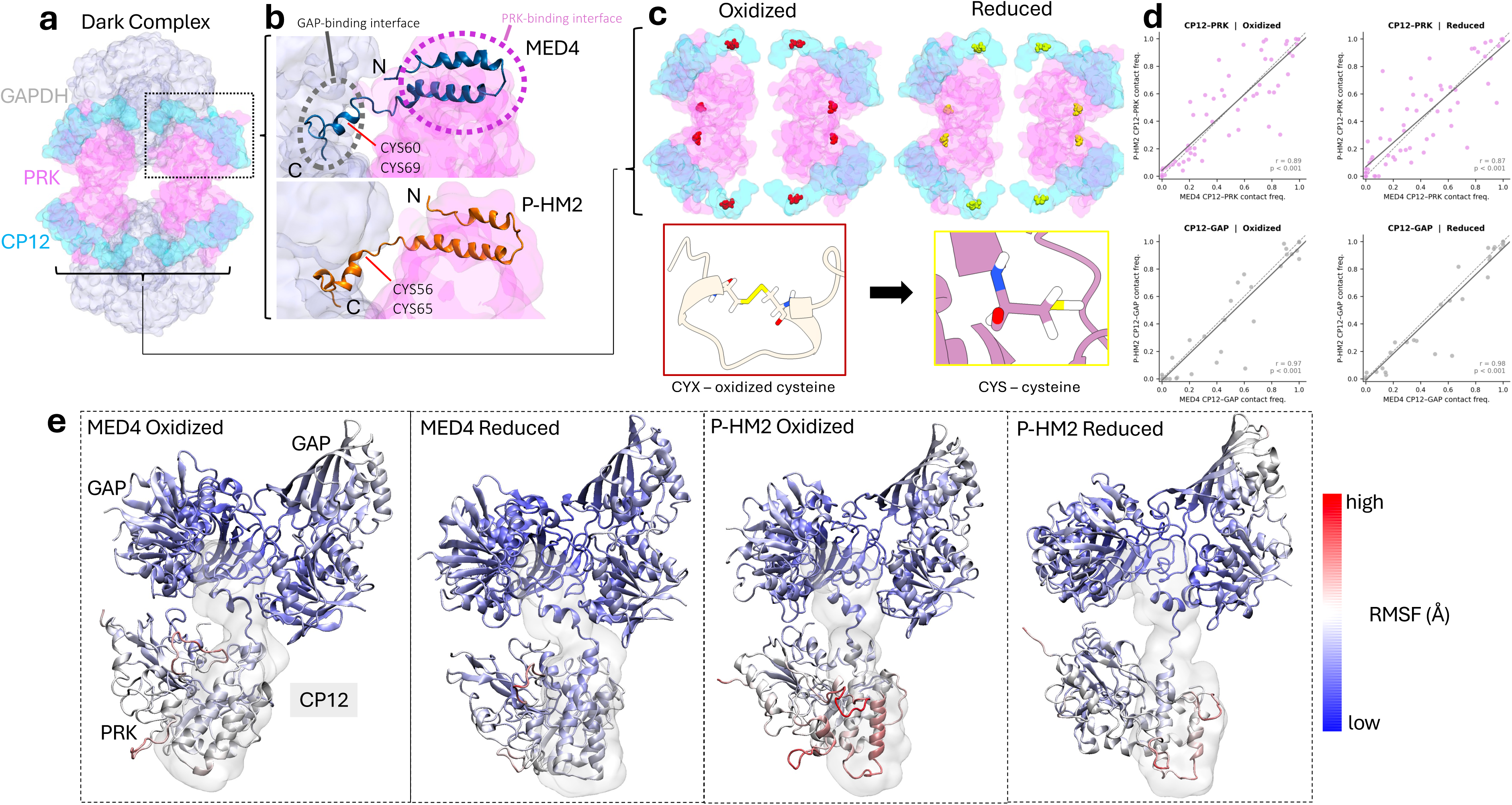
The Dark Complex model, Host MED4 and Viral P-HM2 CP12, Differences between Oxidized and Reduced conditions. **a** The PDB structure of the GAP2-PRK-CP12 complex, GAP2 is colored in grey, PRK in pink, and CP12 in light cyan. **b** Localization of the GAP2-binding and PRK-binding region on the CP12 chain, in blue is MED4 and in orange P-HM2, showcasing each variant. **c** We define the structural differences between the oxidized system and reduced system over the dark complex which is identical for each variant. In the oxidized system, oxidized cysteines (CYX) are placed on residues 60 and 69 of all MED4 CP12 chains, and on residues 56 and 65 of all P-HM2 CP12 chains, and residues 201 and 225 of all PRK chains for both variants. In the reduced system cysteines (CYS) are placed on these same residues. **d** The contact frequency correlation between MED4 and P-HM2 for PRK (top) and GAP (bottom) subunits across each condition, oxidized (left) and reduced (right). **e** The variable Root Mean Squared Fluctuation (RMSF Å) of a single GAP2-PRK-CP12 section of the dark complex (GAP2 chain E, F. CP12 chain C. PRK chain B.) between MED4 (left) and P-HM2 (right), and across each oxidized and reduced condition. Calculated across three replicate molecular dynamics trajectories. CP12 for each system is outlined lightly in grey.

We simulated complexes containing MED4 or P-HM2 CP12 under oxidizing conditions, with intramolecular disulfides in CP12 (MED4 Cys60-Cys69, P-HM2 Cys56-Cys65) and PRK (Cys201-Cys225) (**Fig. 2a-c**), and under reducing conditions with these residues as free thiols (**Fig. 2c**). The CP12-bound fraction remained 1.00 across both variants, fully reduced (“reduced”) and reference states (“oxidized”), and replicates (n=3). CP12 engaged PRK and GAP2 through spatially distinct interfaces, with PRK contacts enriched in the N-terminal and central regions and GAP2 contacts in the C-terminal region (**Fig. 2a-b**). Contact maps were highly correlated between complexes, with Pearson r of 0.87-0.89 at the PRK interface and 0.97-0.98 at GAP2 (**Fig. 2d**). P-HM2 CP12 therefore preserves the host-like binding topology to engage both MED4 PRK and GAP2.

Per-residue root mean square fluctuation (RMSF) after equilibration was consistently higher for P-HM2 CP12 than for MED4, with the largest differences near the PRK-binding region (**Fig. 2e**), though this reached significance only under reduced conditions (p = 0.01; **Fig. S6**). GAP2-binding residues showed smaller, non-significant differences under both states. These results suggest that P-HM2 CP12 may preferentially increase conformational sampling near the PRK-facing interface while preserving the conserved GAP2-facing region. As a follow-on we used Molecular Mechanics/Generalized Born Surface Area (MM/GBSA) binding-energy calculating global van der Waals (VdW) across paired systems, and fully reduced versus reference states (reference is labelled as “oxidized”) these results aligned with the above analysis of differential subunits; P-HM2 CP12 bound PRK more favorably in reduced conditions while in MED4, CP12-GAP2 binding was favored (**Fig. S7**).

### P-HM2 CP12 strengthens PRK interface burial while loosening the GAP2-facing interface by redistributing local VdW interactions through altered residue chemistry and contact dynamics

We conducted an analysis of contact number, frequency, dwell time, turnover, and mean distance for each CP12-partner interaction (**Table S3, Fig. S8**) and examined these compared with residue chemistry shifts (**Fig. S9**). Next we tested whether contacts identified by a 5 Å heavy-atom cutoff were associated with favorable local packing, using residue-level MM/GBSA VdW decomposition as a position-resolved energetic readout and defining VdW favorability as -VdW. P-HM2-associated gains in VdW favorability were most consistently linked to higher contact frequency, longer dwell time, and shorter distance, strongest at the PRK interface (**Fig. 3a**). In oxidized PRK contacts, VdW favorability was positively associated with frequency and dwell time and negatively with distance, retained under reduced conditions (oxidized: n = 153-195, all p < 3.78 × 10⁻¹⁰; reduced: n = 157-195, all p < 5.25 × 10⁻⁵). Hydrophobic/polar-to-charged positions at the oxidized PRK interface carried strong positive frequency- and dwell-to-VdW coupling (n = 36, both p < 0.001; **Fig. 3b**). P-HM2 substitutions, chemistry shifts, contact dynamics, and local packing therefore appear mechanistically linked, with substitutions reshaping which contacts form and persist.

**Figure 3.**
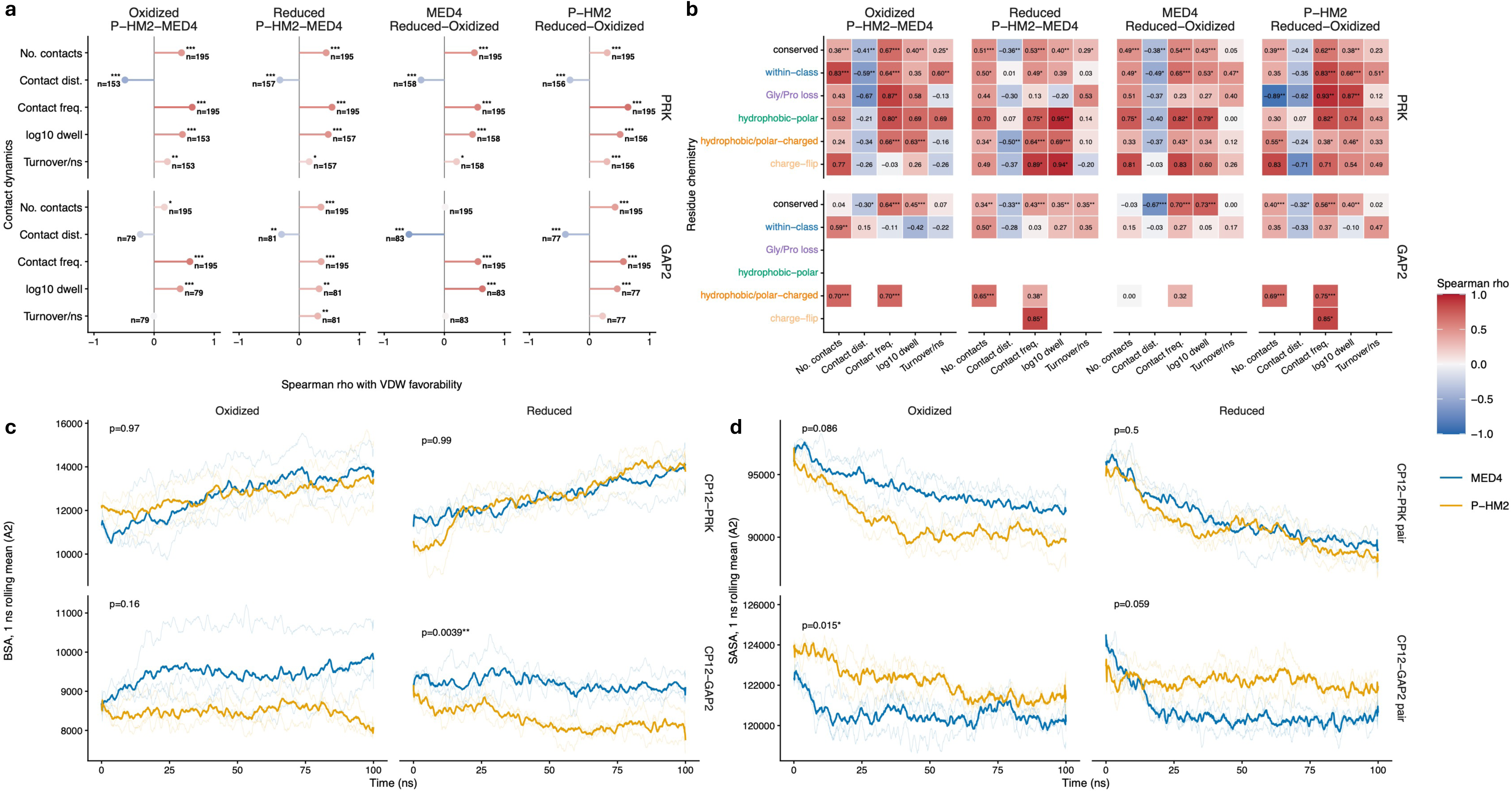
Contact-dynamics, VdW-packaging, and surface-burial signatures of P-HM2 Dark Complex remodeling. **a** Spearman correlations between CP12 contact-dynamics and residue-level van der Waals (VdW) favorability across PRK and GAP2 interfaces, comparing P-HM2 against MED4 Dark Complex under oxidized and reduced conditions, and comparing reduced against oxidized states within each model system. VdW favorability was calculated as the negative MM/GBSA VdW term, so higher values indicate more favorable local VdW packing. Contact dynamics were quantified from 5 Å heavy-atom CP12-partner contacts sampled every 100 ps across 100 ns trajectories and included the number of contacts, mean contact distance, contact frequency, log10 dwell time, and turnover per ns. Positive rho indicates that a larger contact dynamics parameter value is associated with more favorable VdW packing. **b** Residue-chemistry-stratified Spearman correlations for the same comparisons as in a. Tile color shows Spearman’s rho, with positive values indicating stronger coupling to favorable VdW packing and negative values indicating weaker or opposing coupling. To account for multiple testing across the contact dynamics-VdW coupling, Benjamini-Hochberg (BH)-false discovery rate (FDR)-adjusted significance is applied with asterisks indicating p < 0.05 (*), p < 0.01 (**), and p < 0.001 (***). **c** and **d** Time-resolved buried surface area (BSA) and solvent-accessible surface area (SASA) profiles compare MED4 (blue line) and P-HM2 (orange line) Dark Complex simulations under oxidized and reduced conditions. Thin lines represent individual replicate trajectories, and thick lines represent the system-level mean. Columns show oxidized and reduced simulations; rows show CP12 interaction PRK, or GAP2. P values from paired t-test comparing MED4 and P-HM2 replicate-level means within each redox state; asterisks indicate p < 0.05. “Oxidized”, containing disulfide bonds on PRK and CP12 cysteines, served as the reference system in each ensemble. “Reduced”, contain fully reduced cysteines in the complex.

This relationship was also modulated by redox state, with different partner emphasis. In MED4, redox-associated VdW changes were strongest at GAP2, defined by higher frequency, longer dwell time, and shorter distance (n = 83-195, all p < 8.32 × 10⁻⁹; **Fig. 3a**). In P-HM2 they were strongest at PRK, with higher frequency, longer dwell time, shorter distance, and higher turnover (n = 156-195, all p < 2.48 × 10⁻⁴). Chemical-class stratification showed that this coupling was organized differently. In MED4, the strongest GAP2 signal was enriched at conserved positions, whereas P-HM2 PRK coupling extended across substituted classes, including within-class, Gly/Pro-loss, and hydrophobic/polar-to-charged positions (**Fig. 3b**). MED4 redox-sensitive packing may therefore be anchored mainly in conserved GAP2-facing contacts, whereas P-HM2 redistributes it toward PRK-facing substituted positions.

To test whether contact-level remodeling propagated into interface-scale reorganization, we quantified buried and solvent-accessible surface area (BSA, SASA) using the same trajectories. Time-resolved BSA showed P-HM2 did not globally reduce CP12 engagement (**Fig. S10**), and CP12-PRK BSA was nearly unchanged under both states (**Fig. 3c**). The main difference emerged at the CP12-GAP2 interface, where P-HM2 showed lower BSA than MED4, reaching significance under reduced conditions (p = 0.0039), with the same trend under oxidized conditions (p = 0.16, **Fig. 3c**). Complementarily, CP12-GAP2 SASA was higher in P-HM2, significant under oxidized conditions (p = 0.015) and trending under reduced conditions (p = 0.059, **Fig. 3d**). P-HM2 therefore preserves overall engagement while selectively reducing burial and increasing exposure at the GAP2-facing interface, potentially altering accessibility of redox-sensitive residues.

### MED4 redox proteomics reveals distributed cysteine oxidation across the dark complex

These MD simulations implied that substitution-driven interface changes could interact with redox-sensitive host residues during dark complex regulation. We therefore performed MED4 redox-proteomics experiments under illumination perturbation and analyzed the dataset using our established human-in-the-loop agentic AI workflow to help identify oxidation-sensitive cysteines in CP12, GAP2, and PRK with human review, grounding the sites examined in subsequent combinatorial PTM simulations (**Fig. 4a**).

**Figure 4.**
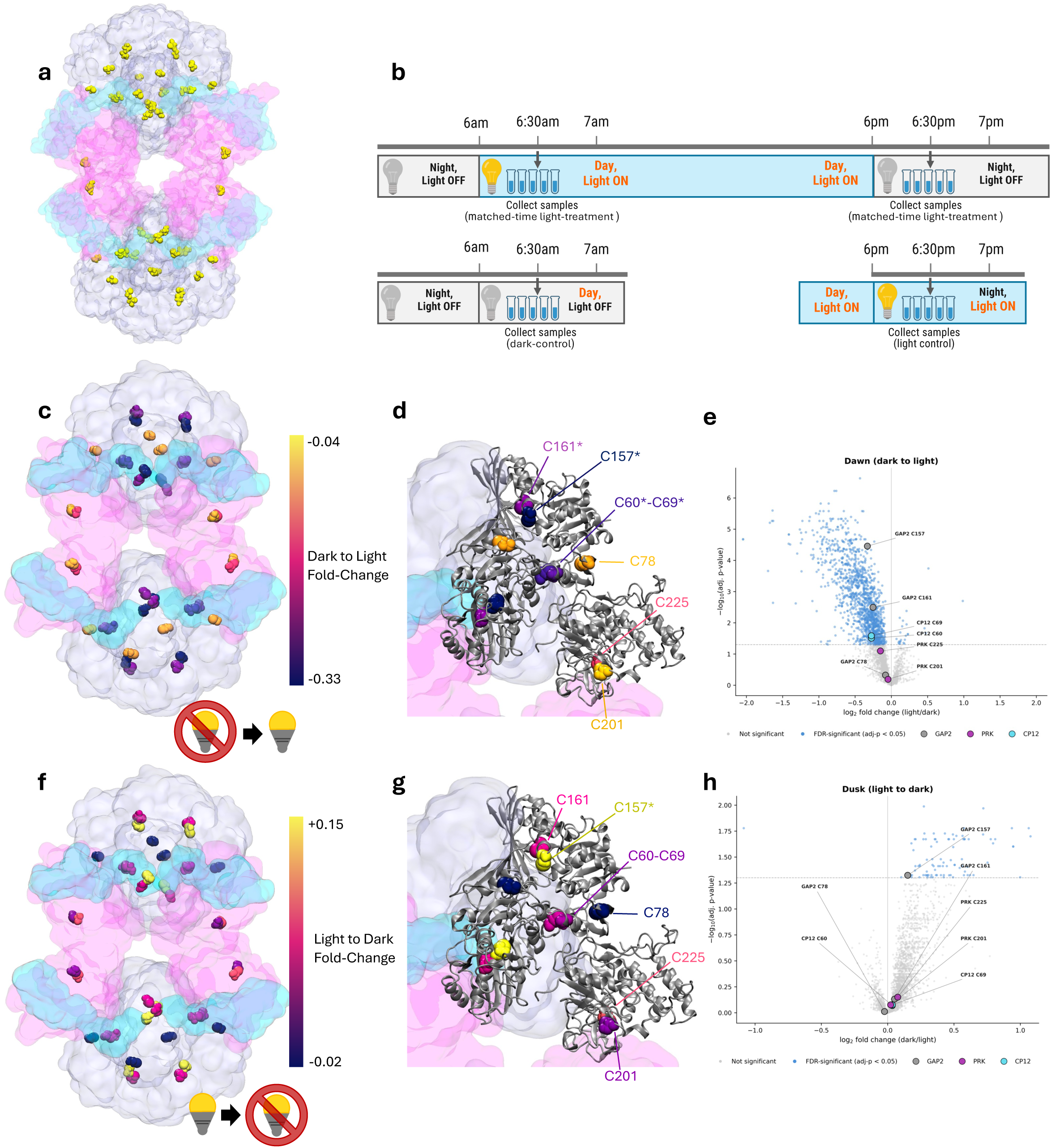
Redox proteomics measurements of the MED4 Dark Complex. **a** Marked model with all cysteines (48 distinct residues) within the dark complex model, which forms the set of possible locations for oxidative modifications, colored in yellow. **b** Experimental design for illustrating the relative fold-change measurement scheme for each scenario (dark-light perturbation and light-dark perturbation). For each distinct perturbation (dawn and dusk) samples are measured relative to control cultures that have been acclimated to dark and light respectively. Samples are measured at 6:30am and 6:30pm. **c** Mapping of experimentally measured fold change values to their respective Cysteine location within the dark complex under illumination disturbance. This coloring represents the dark-to-light perturbation, akin to the state the complex would be in when it becomes reduced. **d** A close-up of one quadrant of the dark complex with two GAP2 chains, one CP12 chain, and one PRK chain. C201 of the PRK chain is the least responsive, while C60 and 69 of the MED4 CP12 chain are moderately responsive. C225 of PRK and the cysteines on GAP2 show the highest fold change, as a measured response to the dark to light shift. **e** Proteome-wide volcano plots of the perturbation contrast across all 3,571 cysteine oxidation PTM sites detected in MED4 by proteomics. The dawn perturbation (dark to light; light treatment vs. matched-time dark control). Each point is one cysteine oxidation PTM site; x-axis shows abundance-corrected log_2_ fold-change (light vs. dark) in oxidation occupancy;, y-axis shows -log_10_(adj. p-value). Translucent points are non-significant (p > 0.05) and blue points are significant (adj. p < 0.05). Specific points pertaining to the dark complex are colored with CP12 in salmon, GAP2 in grey, and PRK in Pink. **f** Experimentally measured fold-change mapped to the dark complex structure for the light-to-dark perturbation. **g** Corresponding close-up of the same dark complex quadrant under light-to-dark perturbation GAP2 CYS157 shows the largest fold-change. **h** Proteome-wide volcano plots for all cysteine oxidation PTM-sites (corrected for protein abundance changes), for the light-to-dark perturbation. Dark complex cysteine changes were not accompanied by significant changes in the relative abundance of GAP2, PRK, and CP12 (Fig. S11), supporting the hypothesis of light-dependent regulation primarily through cysteine redox state change.

The two MED4 CP12 redox-switch cysteines, Cys60 and Cys69 (HMM positions 57 and 66 in P-HM2), showed significantly higher oxidation in the dark-adapted baseline than after dawn illumination (p = 0.031), consistent with redox cycling during the light perturbation (**Fig. 4b**). Oxidation was detected at five GAP2 and PRK positions. Among these sites, only GAP2 Cys157 and Cys161 showed a statistically significant change under light perturbation (Cys157, log_2_FC=-0.330, adj-p=3.50e-5; Cys161, log_2_FC=-0.253, adj-p=0.0032), whereas PRK Cys225 showed a nominal trend and GAP2 Cys78 and PRK Cys201 were not significant (**Fig. 4c-d, Fig. S11**). Under dark perturbation, the redox response was less extreme at the 30 min post-perturbation timepoint; only GAP2 Cys157 showed significant change (**Fig. 4f-g**). In both perturbations, these cysteine changes were not accompanied by significant changes in the relative abundance of GAP2, PRK, and CP12 (Fig. S11), supporting the hypothesis that light-dependent regulation acts primarily through cysteine redox state change. These measurements support a secondary hypothesis that dark complex redox regulation arises from the combined modification states of CP12 and its partners rather than CP12 disulfide switching alone, grounding the sites sampled in the PTM simulations below. In dark complex components, we observed a stronger redox response to the dawn (dark-to-light) perturbation than to the dusk (light-to-dark) perturbation; this follows a general trend across the proteome. After correcting for protein abundance changes, the number of significantly changed Cys redox intensities and the average absolute fold change of those intensities are both substantially greater in response to the introduction of light at dawn (**Fig. 4e**) than in response to the removal of light at dusk (**Fig. 4h**).

### Combinatorial PTM modeling identifies GAP2-centered conformational divergence

We next asked how multiple, potentially coordinated oxidative cysteine modifications alter the stability and dynamics of the host and phage dark complexes. For each model system, we generated 1,011 additional combinatorially PTM-configured structures covering the cysteines in the dark complexes, yielding 1,012 systems including the unmodified reference (referred to as “REF”) (**Fig. 5a**) that match the chemical space of the dark complex from the redox proteome. Selected GAP2 and PRK cysteines were assigned *S*-nitrosylation (SNO), *S*-sulfenylation (SOH), or *S*-glutathionylation (SSG), while CP12 and PRK were also modeled with intact or reduced disulfide bonds; complete assignments are provided in **Supplementary Tables S4 and S5**. Each system underwent a 100 ns all-atom MD simulation with explicit solvent models (∼close to 800k atoms) followed by structural and dynamical analyses. Across the 1,012 systems in each ensemble, simulations showed substantial system-to-system variability in RMSD, RMSF, radius of gyration (Rg), and secondary-structure composition (**Fig. S12**). We utilized principal component analysis (PCA), composed of eight structural descriptors: mean backbone dihedral angles (φ and ψ), B-factors, coil, helix, and sheet fractions, RMSD, and Rg, to rank the systems by variance-normalized Euclidean distance to the reference (REF). We found PTM burden was associated with greater conformational divergence (**Fig. S13a**) and PCA distance was correlated mainly to GAP2 thiol modification (**Fig. S13b**). GAP2 PTM number correlated strongly and positively with PCA distance in both variants (Spearman ρ = 0.68 in MED4, 0.72 in P-HM2). PTM counts for PRK and CP12 were weakly correlated (**Fig. S13b**). This indicates cysteine PTMs on GAP2 may modulate redox-induced conformational divergence.

**Figure 5.**
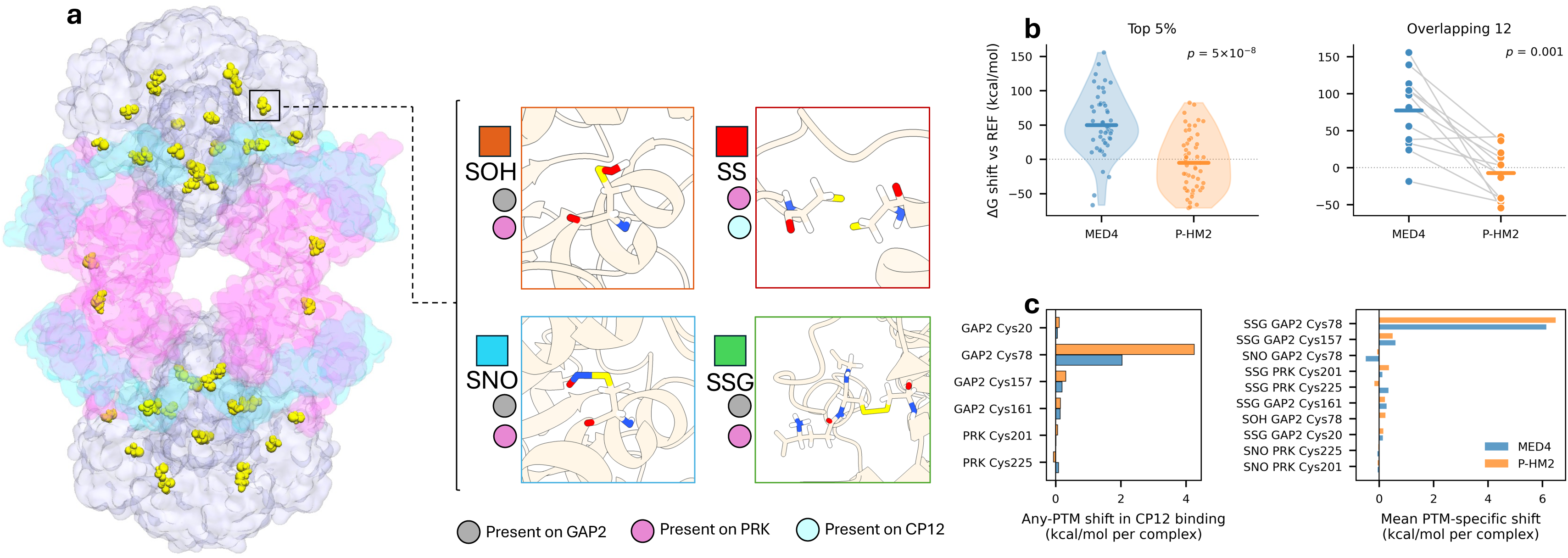
Dark Complex combinatorial PTM simulations. **a** The dark complex system is modified combinatorially for both variants, with four possible PTM types: a disulfide (SS), S-nitrosylation (SNO), sulfenylation (SOH), or glutathionylation (SSG). For any given CYS site on either variant complex GAP2 is modified by only SNO, SOH, and SSG at residues 20, 78, 157, and 161; PRK by any type at residues 201 and 225; CP12 only SS at residues 60 and 69 for MED4 and residues 56 and 65 for P-HM2. **b** Metrics from all 2,024 all-atomistic molecular dynamics simulations of MED4 and P-HM2 modified dark-complexes. MED4 (left) and P-HM2 (right) from top to bottom the Root Mean Squared Deviation (RMSD) (Å), Root Mean Squared Fluctuation (Å) compared to the oxidized reference system, Radius of Gyration (Rg) (Å), and the secondary structural composition are tabulated. For RMSD and R_g_ the reference system is shown in black.

Comparative binding free energy calculations provide a high-resolution *in silico* follow-up to sites identified by redox proteomics^24^. Using the variance-normalized ranked simulations, we computed MM/GBSA binding-energy shift (ΔG_shift, vs. REF) of CP12 binding with GAP2 and PRK under PTMs compared to their respective unmodified references. Comparing MED4 and P-HM2 sets (**Fig. 5b, left**), PTMs significantly weakened CP12 binding in MED4 but left it essentially unchanged in P-HM2. This was clearest among the 12 systems shared between both top 5% selections, which carry identical GAP2 and PRK PTMs and therefore isolate the effect of CP12 origin. In 11 of 12 matched pairs, the same PTM combination weakened CP12 binding more in MED4 than P-HM2 (**Fig. 5b, right**); the exception was system 0620 with GAP2 Cys157 sulfenylation. P-HM2 CP12 therefore buffers the binding-energy costs imposed by identical modifications on the host complex.

Per-residue decomposition localized this cost to GAP2 Cys78, whose contribution shifted by +2.0 kcal/mol in MED4 and +4.2 kcal/mol in P-HM2, the largest among all sites; remaining cysteines each contributed less than 0.3 kcal/mol (**Fig. 5c, left**). This was driven predominantly by glutathionylation (**Fig. S14**), which shifted Cys78’s contribution by +6.2 (MED4) and +6.5 (P-HM2) kcal/mol, roughly an order of magnitude above the next site-chemistry combination (**Fig. 5c, right**). The unfavorable local contribution of Cys78 glutathionylation was thus similar in both systems, despite minimal net change in P-HM2 CP12 binding.

### Conformational flexibility enables P-HM2 CP12 to accelerate cluster kinetics and buffer PTM-associated binding costs

We next asked how P-HM2 CP12 buffers the binding-energy cost destabilizing MED4. Partner-partitioned binding energies showed the MED4 penalty localized mainly to the PRK-facing interface (+32 kcal/mol, paired p = 0.009; top 5%, +46 kcal/mol, p = 3 × 10⁻¹³) and CP12-associated interaction terms (+147 kcal/mol, p = 5 × 10⁻⁴), whereas P-HM2 terms were unchanged or modestly stabilized (**Fig. 6a**). GAP2-side changes were similar between systems, indicating host GAP2 contact loss was energetically compensated whereas PRK-side loss was not. Matched perturbations produced opposite dynamical responses: MED4 contracted and rigidified, with decreased Rg across all subunits and reduced mobility near PRK, whereas P-HM2 expanded and remained mobile (whole-complex ΔRg, −1.6 Å vs. +2.0 Å, paired p = 5 × 10⁻⁴; PRK ΔRMSD, −1.6 vs. +1.3 Å, p = 0.007; **Fig. 6b-c**). This Rg trend held across the top 5% ensembles (whole-complex, p = 4 × 10⁻¹⁴; PRK, p = 4 × 10⁻¹⁶). Consistent with these opposing dynamical responses, kinetic network analysis showed that mean first-passage time (MFPT) varied across matched PTM states, whereas intra-cluster residence time (τ) was generally prolonged in MED4 and shortened in P-HM2 relative to their respective reference states, indicating more frequent escape from locally clustered conformational states in the P-HM2 system (**Fig. S15**). MED4 also showed a larger reduction in buried CP12 surface fraction (−0.047 vs. −0.011, p = 0.001) and lost contacts with both partners (GAP2 per residue, −0.032 vs. +0.009, p = 0.001; PRK, −0.172 vs. +0.003, p = 0.034; **Fig. 6d–f**, see supplementary Movie.S1), reproduced across the top 5%. P-HM2 CP12 therefore buffers the GAP2 Cys78 perturbation by remaining expanded, mobile, and engaged across the interface

**Figure 6.**
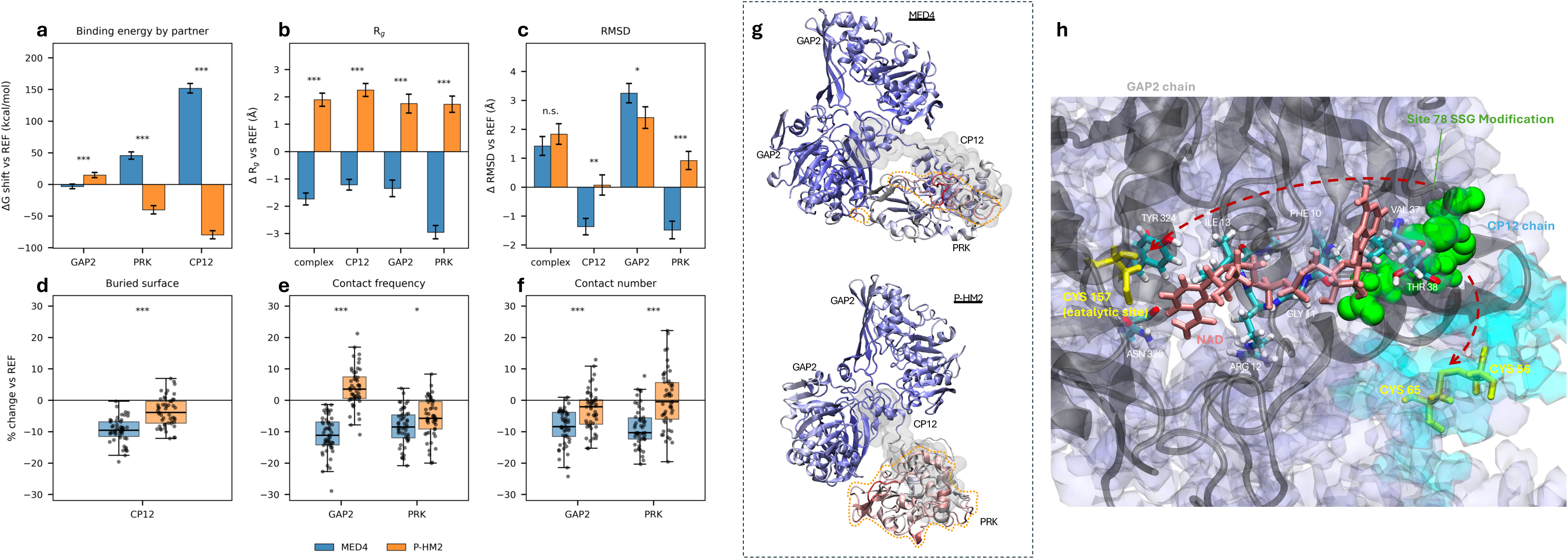
Conformational flexibility of P-HM2 CP12. Matched overlap-12 systems shown. MED4 blue, P-HM2 orange. Top-row panels (**a-c**) are bars (mean ± SEM); bottom-row panels (**d-f**) are box plots (center line, median; box, IQR; whiskers, 1.5 IQR) with individual systems overlaid as points. Significance is two-sided paired Wilcoxon throughout, asterisks denote statistical significance (* p<0.05, ** p<0.01, *** p<0.001). All quantities are change relative to each ensemble’s own unmodified reference (Δ vs REF). Top row, energetic and conformational response. **a** Binding-energy change partitioned by receptor partner, **b** change in radius of gyration, and **c** Cα RMSD by subunit. Bottom row, interface response as percent change vs REF. **d** Fraction of CP12 surface buried, **e** contact frequency per CP12 residue, and **f** number of contact pairs per CP12 residue to GAP2 and PRK. **g** A GAP2-PRK-CP12 quadrant of the dark-complex system 0928 (the most different of the overlapping systems) colored by RMSF. MED4 (top) shows small regions of high flexibility in PRK proximal to the CP12 binding region, while P-HM2 (bottom) shows a large region of high flexibility widespread in PRK, both high RMSF regions are marked by dashed yellow lines. **h** A close up of GAP2 chain A, and CP12 chain P, on P-HM2 system 0705. The GAP2 catalytic site C157 is highlighted alongside the NAD cofactor (highlighted in pink), GAP2 Glutathione modification (colored in green) is localized to position 78 of GAP2 in proximity to the viral CP12 disulfide bonded residues 56 and 65. GAP2 C78 is located such to facilitate the propagation of changes in local packing to both the distant catalytic site C157, across the labelled intermediate residues, and the nearby CP12 chain.

## Discussion

Cyanophages shape marine primary production globally^7^, yet how their auxiliary genes manipulate host carbon metabolism remains unclear. Viral auxiliary metabolic genes encoding enzymes reroute host flux directly^11,25^, whereas viral regulatory proteins must engage existing host complexes while altering their assembly, environmental responsiveness, and enzymatic regulation. CP12 is well suited to probe this because it is both a key regulator of host carbon fixation that has also been detected in phages and an intrinsically disordered protein whose function depends on cysteine control, conformational dynamics, and transient interface formation, none of which can be resolved by static structural prediction alone^5,26,27^. We therefore connected protein-family-wide CP12 analysis with paired host and phage MD simulations, constrained by redox proteomics in *P. marinus* MED4. Our analysis suggests cyanophage P-HM2 CP12 preserves structural compatibility with host GAP2 and PRK while remodeling the physicochemical, contact-dynamic, energetic, and redox-sensitive features of partner engagement.

Viral CP12 appears to achieve host compatibility through physicochemical rather than sequence mimicry. At the slow evolutionary scale of host adaptation reflected in the protein-family analysis, CP12 shows high residue-level entropy but strong conservation of chemical class and cysteine placement, so identities can vary while key chemical properties are maintained. This agrees with prior observations that protein evolution preserves physicochemical interaction properties despite changes in identity, and with cyanophage auxiliary proteins retaining host-like functions despite divergence^11,28,29^. Our comparison extends this by showing that viral CP12s, predominantly in tailed double-stranded DNA phages, remodel the chemistry of the interface positions that is more often hydrophobic or polar in bacterial CP12, with higher enrichment of charged features and selected gains in polar or Gly/Pro geometry-modulating features. Such a shift could reduce dependence on surface complementarity, preserving host protein compatibility while increasing docking flexibility and reversibility on a molecular dynamics scale^30^, consistent with viral imitation of host-like function without full sequence similarity ^11,29,31^. We therefore hypothesized that viral CP12 retains the compatibility required to engage the host dark complex, while N- and C-terminal substitutions differentially affect GAP2 and PRK interactions. Supporting this, the paired simulations showed P-HM2 CP12 preserves dark-complex association while increasing PRK-facing contact persistence and van der Waals interactions yet reduces GAP2-facing burial and leaves that surface more solvent-exposed.

This partner-specific remodeling offers a molecular basis for cyanophage control of host light-dependent carbon metabolism. An enzymatically inactive dark complex assembly begins with CP12 binding GAP2, followed by PRK recruitment driven by the redox PTMs of cysteines^11,27,29^. Oxidized CP12 folds into a hairpin-like conformation that binds both GAP2 and PRK into a dark complex, thereby burying and blocking access to their binding sites. When photosynthetic processes take place, reduced CP12 is unstructured and dissociates from its partners. Enzymatic activities of GAP2 and PRK resume when their metabolite binding sites are exposed again. Conservation of the GAP2-facing interface may preserve the initial assembly step, whereas divergence at the PRK-facing interface could tune the second, including PRK recruitment, sequestration, or release. Because cytosolic PRK availability influences the balance between Calvin-Benson activity and oxidative pentose phosphate activation^32^, viral remodeling of PRK engagement may redirect reducing power and carbon flux during infection. This is consistent, though not specific, with reported co-expression of CP12 alongside photosynthetic light reaction, deoxynucleotide biosynthesis, and pentose phosphate genes, and with altered NADPH/NADP and NADP(H)/NAD(H) ratios in infected cells^11^. Such regulation is biologically relevant because *Prochlorococcus* coordinates carbon fixation, respiratory carbon use, cell-cycle progression, and central-carbon regulation across the diel cycle, while P-HM2 infection is itself light dependent^33–35^. Our recent genome-scale modeling showed P-HM2 CP12 can redirect host carbon flux toward the pentose phosphate pathway and nucleotide metabolism, with nitrogen-dependent growth effects validated in a tractable cyanobacterial system^10^. However, whether cyanophage CP12 physically assembles with host GAP2 and PRK has not been directly established. Our simulations address this step by testing whether a divergent homolog remains compatible with the host architecture while altering partner-specific interactions, providing a testable hypothesis for future biochemical and physiological study to determine whether phage CP12 assembles with host GAP2 and PRK *in vivo* and alters their carbon metabolism during infection.

The same interface asymmetry indicates which residues are prone to redox control, since changes in GAP2 burial and exposure could place redox-sensitive residues where local modifications propagate. Our redox proteomics under illumination disturbance highlights how the post-translationally modified redox state of cysteine residues can change bidirectionally between light and dark phases of the day, independent of protein abundance. We identified coordinated cysteine oxidation across CP12 and GAP2, including GAP2’s catalytic nucleophile site Cys157 and nearby Cys161, showing the strongest response, while the changes in the relative protein abundance remain insignificant. This finding is consistent with cysteine oxidation acting as a rapid, reversible mechanism coupling protein structure and interaction state, through *S*-nitrosylation, *S*-sulfenylation, *S*-glutathionylation, and disulfide exchange^32,33^ in response to cellular redox conditions, in contrast to slow protein expression. The MED4 redox and global proteomic datasets defined the experimentally grounded cysteine-modification landscape for constructing the PTM simulations. The thiol PTM molecular dynamics simulations quantified this pattern mechanistically, with GAP2 PTM load producing the largest conformational shifts.

Modification of <u>GAP2</u> Cys78 by glutathionylation introduces a bulky glutathione group near the GAP2-CP12 interface that disrupts local packing, weakens CP12 binding and propagates conformational changes, resulting in the buried GAP2 catalytic site, Cys157 over distance (**Fig. 6g**). Cys78 was detected in the MED4 redox proteomics dataset, although its regulation did not increase or decrease significantly during either perturbation. Despite this limited evidence for a strong physiological redox transition under the sampled conditions, glutathionylation of Cys78 imposed the highest CP12 binding-energy cost among all tested site and PTM combinations in the simulations. This identifies Cys78 as a potentially high-impact regulatory site when modified, consistent with prior evidence that GAP2 can undergo reversible glutathionylation in plants and cyanobacteria^36,37^. Furthermore, our recent PTM-Psi studies identified Cys78 as redox-sensitive near cofactor-binding sites in homolog models^17,38^. Although Cys78 is distal to the catalytic site, its modification has been reported to perturb NADP+ positioning through steric interference with the adenosine 2′-phosphate group, disruption of Arg80-associated cofactor positioning, and loss of a stabilizing thiol-phosphate interaction^17,38^. Because glutathionylation forms a mixed disulfide that is reversible but generally requires enzymatic deglutathionylation^40^, this modification may be kinetically persistent under oxidative or infection-associated redox conditions. Cys78 SSG could therefore act as a redox brake on GAP2 catalytic readiness, prolonging GAP2-mediated inhibition while CP12-mediated PRK engagement maintains the dark-complex state. Both the *in vivo* oxidation and *in silico* conformational impact suggest GAP2 as a candidate site transducing redox state into conformational change, extending established models of CP12-mediated redox regulation^27^. These simulations focused on cysteine oxidation, while other regulatory modifications, including phosphorylation, acetylation, and mixed PTM states, remain important extensions for testing how additional cellular signals intersect with CP12-mediated dark-complex regulation.

We integrate these observations of tuning a broad chemical space into a new variation of the conformational rheostat model^41^, in which P-HM2 CP12 does not lock the complex into a single inhibited state but preserves responsiveness while altering the balance among accessible states. This follows the characteristics of a protein-rheostat framework, in which amino acid substitutions tune functional output through local structural perturbations or changes in dynamic coupling. Three observations support the combinatorial thiol PTMs that finely tune over the structural continuum of a protein rheostat in response to redox homeostasis^42,43^. First, the same PTM configurations did not produce the same degree or direction of structural displacement in MED4 and P-HM2, so PTM identity alone does not determine conformational outcome. Second, across the 12 matched PTM states, P-HM2 CP12 shortened cluster residence time relative to its reference while leaving first-passage time largely unchanged, whereas MED4 CP12 lengthened both, consistent with more frequent escape from local conformational states. Third, increased flexibility of the viral CP12 chain at the PRK interface, with the expanded, mobile complex seen under PTM stress, indicates enhanced sampling rather than rigid stabilization. Because the comparisons are based on 100 ns trajectories across a large set of PTM states, they are best interpreted as relative shifts in accessible conformational dynamics (see Supplementary Movie 2). Slower transitions could be further resolved with longer replicates, enhanced sampling, or Markov-state modeling.

Our study demonstrates a cross-scale framework in which viral CP12 divergence reflects evolutionary adaptation to host regulatory machinery, while its functional consequences emerge through physicochemical interactions operating at faster molecular timescales. At a slow, evolutionary scale, phage CP12 does not appear to gain advantage by globally stabilizing a sturdier dark complex. Instead, divergence at partner-facing interfaces tunes local contact behavior, favoring closer and more persistent PRK engagement while preserving GAP2 compatibility for complex assembly. This provides a first proposed mechanism for viral control, efficient partner capture, and PRK sequestration without a large increase in overall complex stability. A second mechanism emerges under redox and PTM stress at a fast, molecular scale, the redox state of cysteine residues, as reflected by PTM levels, can change bidirectionally between light and dark phases of the day, independent of protein abundance. Thiol PTM mechanistically introduces a strong GAP2-centered perturbation tuned by the redox of CP12 that increases its cost of folding and binding and may reduce GAP2 catalytic readiness, yet the P-HM2 simulated complex remains more expanded, mobile, and contact-rich, maintaining PRK engagement instead of transmitting the perturbation into PRK-interface weakening. Thus, phage CP12 may prolong dark-complex inhibition by combining partner-specific contact redistribution with buffering of redox perturbation. This mechanism could favor Calvin-Benson cycle inhibition and redistribution of reducing power toward the pentose phosphate pathway and nucleotide biosynthesis during infection^5,11,13^. More broadly, our framework connects viral sequence divergence, interface chemistry, molecular dynamics, and redox-sensitive metabolic regulation, providing a basis for understanding how divergent viral regulatory proteins can hijack host pathways by dynamically remodeling existing protein complexes.

## Materials and Methods

### Environmental metagenome assemblies and annotation

Metagenome assemblies were derived from 101 environmental samples collected from the Salish Sea estuary in a prior study^23^. The Salish Sea is a complex water body shaped by highly variable tidal conditions, freshwater outflows, and diverse adjacent landscapes, creating a range of habitats well suited to sampling microbial diversity and capture host-phage adaptation. Surface-water samples were collected across varying times of day and tidal conditions from five areas along the northern Olympic Peninsula, the north inlet of Sequim Bay (a floating dock, where most samples were collected), the south end of Sequim Bay, Cline Spit, Discovery Bay, and north of Port Townsend, from March through October 2024, with a heavy focus on summer months. DNA was extracted and sequenced by Azenta Life Sciences (South Plainfield, NJ, USA).

Contigs assembled in the previous study^23^ were filtered to ≥1 kb, and protein-coding sequences were predicted with Prodigal (v2.6.3) in metagenomic mode. Predicted proteins were annotated with KEGG Orthology using KofamScan (v1.3.0; E ≤ 10⁻³) and with Pfam-A (v37.0) domains using HMMER hmmscan (v3.3.2) under trusted cutoffs. Contig-level taxonomy was assigned with the Contig Annotation Tool (CAT, v6.0), classifying filtered contigs independently against GTDB (2023-11-21) and NCBI NR (2024-04-22); NR lineages were reformatted into GTDB-style rank-prefixed strings for integration. Viral contigs were identified with geNomad (v1.10.0) on contigs ≥10 kb, and per-sample viral summaries were retained as an additional evidence layer during the annotation merge.

### CP12 candidate identification from the metagenome and reference database

Candidate CP12 proteins were retained when they carried the PF02672 CP12 domain and were assigned to Viruses, Bacteria, Archaea, Fungi, Viridiplantae, Metazoa, or SAR, using a greedy strategy that accepted the broadest available taxonomic evidence whereas entries without resolvable support were excluded. Sequences were retrieved from per-sample protein files, and candidates were split into full-length and partial subsets by gene-call completeness. Public references were retrieved via an automated retrieval script using NCBI Entrez Direct utilities (esearch, efetch, xtract) and the UniProt REST API (accessed 24 Dec 2025). Two curated RefSeq entries were retrieved by accession (P-HM2 CP12, YP_004323596.1; MED4 CP12, WP_019478051.1); additional RefSeq CP12 proteins were retrieved with the query “CP12[Title] AND 40:120[SLEN] AND srcdb_refseq[PROP]”, and reviewed UniProt entries with “(xref:pfam-PF02672) AND (reviewed:true)”. Organism, TaxID, and lineage were stored per sequence, with UniProt lineages normalized to NCBI taxonomy. All sources were pooled with the metagenome-derived sequences and dereplicated at 100% amino acid identity with CD-HIT (v4.8.1; -c 1.0 -n 5), retaining one representative per cluster. Dereplicated sequences were then screened with biochemical and architectural criteria to reduce false-positive PF02672 assignments, removing sequences <50 or >250 amino acids. The resulting curated, dereplicated set was used for all downstream diversity, architecture, interface, and classification analyses.

### HMM-based coordinate mapping and structure-based CP12 interface annotation

Curated sequences were aligned to the PF02672 profile HMM with HMMER hmmalign (v3.3.2). Wrapped Stockholm alignment blocks were concatenated per sequence, and match-state columns were identified from the reference annotation line (#=GC RF) as positions where the RF character was neither “.” nor “-”, then converted to one-indexed HMM coordinates. This defined a shared coordinate system spanning metagenome sequences, references, structural models, interface annotations, per-column composition tables, and classifier features. CP12 interface residues were defined from the static MED4 and P-HM2 dark-complex structures (frame-0 solute-only PTM-Psi systems, preserving the residues, cofactors, and chain assignments used in the trajectory analyses). Chains were mapped to CP12, PRK, and GAP2, and all heavy-atom pairwise distances were computed. A CP12 residue was called an interface residue when any heavy-atom pair fell within 5.0 Å of PRK or GAP2. Residue-level interface annotations recorded CP12 contact with PRK, GAP2, or both. Structural CP12 residues were projected onto the PF02672 coordinate system, transferred to HMM columns, and collapsed across CP12 chains to yield system-specific per-column interface annotations for MED4 and P-HM2, which were then compared in the shared coordinate system.

### Residue-level conservation and composition analyses

Amino acid counts at each PF02672 match-state column were computed across the full curated dataset and within sequence groups, and used to derive alignment occupancy, cysteine fraction, Shannon entropy, residue identity dominance, and chemical-class dominance. Alignment occupancy was the fraction of sequences bearing a non-gap character, and cysteine fraction the number of cysteines divided by the non-gap count at that column. Shannon entropy was computed per column by converting counts of the 20 standard amino acids to frequencies, multiplying each non-zero frequency by its base-2 logarithm, and negating the sum; gaps were tracked separately and excluded from the denominator. Residue identity dominance was the maximum single amino acid fraction, corresponding to the proportion of non-gap residues held by the most common residue. Chemical-class dominance was the maximum class fraction per column, where residues were grouped into six classes: cysteine (C); positive (K, R, H); negative (D, E); polar (S, T, N, Q); hydrophobic (A, V, I, L, M, F, W, Y); and Gly/Pro (G, P). These metrics were merged with the interface annotations, and cysteine-enriched columns were annotated together with neighboring interface columns to test whether conserved cysteines fell within or adjacent to contact positions. For lineage-level analysis, lineages with < 3 sequences were grouped as “Other,” and columns below the 10th-percentile non-gap count within a lineage were filtered for insufficient support; per-lineage, per-column, per-interface-class amino acid fractions were then computed to distinguish whether positions were conserved through specific amino acid identity or broader chemical-class preservation. A permutation null model tested whether observed chemical class dominance exceeded arbitrary class groupings (**Supplementary Information**).

### Physicochemical property analysis

For each substitution between HMM-aligned MED4 and P-HM2 CP12 sequences, charge change was the net charge of the P-HM2 residue minus the MED4 residue at pH 7.0 (K/R/H = +1/+1/+0.5; D/E = −1; else 0), hydrophobicity change the Kyte-Doolittle difference (−4.5 to +4.5), and volume change the difference in van der Waals volumes (Å³).

### Interpretable supervised classification of viral and bacterial CP12

A per-sequence, per-position feature table was built from the HMM-aligned dataset (Python v3.9.17; pandas v2.2.2; NumPy v1.23.0), recording aligned residue state, gap status, lineage, source, and chemical class at each column. Sequences in bacterial or viral domain categories were retained, and categorical features were one-hot encoded (scikit-learn v1.6.1, handle_unknown=“ignore”), encoding each observed category at each HMM position as a binary indicator while ignoring unseen categories during prediction. Because the encoded residue and chemical-class features were sparse, binary logistic regression was used as a transparent classifier whose coefficients map back to HMM positions and chemical-class states (N_viral = 49; N_bacterial = 1,316), with L2 regularization (C = 1.0), the LIBLINEAR solver, balanced class weights, and max_iter = 2,000. Performance was evaluated by stratified 5-fold cross-validation (shuffled, seed 1), reporting fold-level ROC AUC, average precision, accuracy, and F1; out-of-fold viral probabilities generated pooled ROC and precision-recall curves and labels at a 0.5 threshold, with pointwise 95% confidence intervals from 500 bootstrap replicates (seed 1). A final model fit on the full dataset provided coefficients for feature interpretation; because these were estimated from the full-fit model rather than out-of-fold models, coefficient magnitudes were used for feature ranking. Position-level enrichment tests were performed in R (v4.5.1): per-column Fisher’s exact tests (Monte Carlo p-values, B = 10,000, seed 1; gaps and unassigned residues excluded) identified columns with viral-versus-bacterial chemical-class differences, and class-level Fisher tests were adjusted by Benjamini-Hochberg FDR (cutoff 0.01).

### MED4 cell culture and sampling

*Prochlorococcus marinus* strain MED4 (CCMP1986; NCMA, East Boothbay, ME) was propagated in Pro99 medium based on Salish Sea seawater^44^ with nitrogen and phosphorus at twice standard Pro99 concentrations, at 22 °C under a 12 h light:12 h dark cycle transitioning sharply at 06:00 and 18:00 (∼45 µmol photons m⁻² s⁻¹ PAR; LI-250A quantum meter, LI-COR). Cultures were acclimated to this diel rhythm for ≥6 months to ensure a stable circadian rhythm, grown in air-sparged 2 L glass vessels (∼1.6 L medium) to mid-to-late exponential phase (∼0.5–1 × 10⁸ cells/mL; OD₇₅₀ calibrated against flow cytometry^43^). Aliquots of 25 mL were dispensed into vented 50 mL bioreaction tubes (CELLTREAT) and returned to the incubator under the standard cycle. At 18:00 the day before collection, five tubes were foil-wrapped to remain shielded through the upcoming dawn and retrieved at 06:00 just before lights-on (dark-adapted, n = 5); five unwrapped tubes were harvested 30 min later at 06:30 after lights-on (light-exposed, n = 5). All dark-condition handling was carried out in a darkened cold room illuminated only by dim green light. Tubes were centrifuged (8,000 × g, 6 min, 4 °C) and the supernatant discarded; pellets were resuspended in ice-cold PBS (pH 7.4), transferred to microcentrifuge tubes, re-centrifuged under the same conditions, snap-frozen in liquid nitrogen, and stored at −80 °C.

### LC-MS/MS analysis of MED4 protein thiol oxidation

Thiol oxidation along with protein abundance was profiled at 6:30am and 6:30pm in two light conditions (lights-on, or lights-off, as described in the previous section) at each of the two timepoints (n=5 replicates). The 6:30am and 6:30pm timepoints correspond to 30 minutes after subjective dawn or dusk, respectively. Profiling was performed using an automated single-pot, solid-phase-enhanced (SP3) PTM workflow^45^. Briefly, frozen pellets were lysed in ice-cold buffer (250 mM MES pH 6, 5% SDS, 1% Triton X-100, 10 mM EDTA, 100 mM NEM, 10 mM nicotinamide, 10 mM sodium butyrate, 2 mM SAHA), incubated 30 min at room temperature in the dark then 1.5 h at 37 °C with intermittent sonication^17^, and bead-beaten (2 × 30 s, speed 6.5). Per sample, 500 µg protein underwent automated SP3 cleanup and digestion (bead:protein 5:1) on a KingFisher Flex (Thermo), with offline digestion on a ThermoMixer (850 rpm, 37 °C, 1.5 h) using trypsin (Promega) and Lys-C (Wako) at 1:100. Peptides were TMTpro 18-plex labeled (2.5:1) and desalted (C18 Sep-Pak, Waters), then enriched for modified cysteine-containing peptides by Resin-Assisted Capture (RAC)^45–48^, IAM-alkylated, desalted, and high-pH reverse-phase fractionated into 6 fractions^45^. Fractions were analyzed on a Vanquish Neo UHPLC coupled to an Orbitrap Exploris 480 (2 h gradient; DDA, full MS 400–1,800 m/z; top-20 HCD). Spectra were searched with MS-GF+^49^ against the MED4 proteome ^50,51^, processed with PlexedPiper (v0.4.2)^52^, and normalized and analyzed with ProteoMeter (v0.4.0)^53^ to isolate CYS oxidization changes from protein abundance changes. To assist with mapping time-resolved redox abundances onto the dark-complex structures, we developed a repository with selected bioinformatics skills^54^ and specific biological context to ground an AI coding agent running in Visual Studio Code^55^ and Copilot Chat. The AI agent enabled rapid proteomics data exploration and triage of candidate redox-sensitive sites for downstream analysis. To visualize the proteome-wide redox response to the dawn-light perturbation, PTM sites were filtered to cysteine oxidation only (Type = Ox; 3,870 of 13,188 total PTM sites) and retained if intensity data were present in at least one replicate per condition (total missingness ≤ 5 across all six diel conditions), yielding 3,571 sites. For each site, the abundance-corrected log_2_ fold-change and Benjamini-Hochberg (BH) FDR-adjusted p-value for the Light_0630 vs. Dark_0630 contrast (matched-time light-treatment vs. dark-control comparison) were plotted as -log10(adj. p) against log_2_ fold-change; BH correction was computed across the full proteome-wide site set (n ≈ 13,188). Sites were classified as FDR-significant at adj-p < 0.05 (1,439 of 3,571 sites). Seven pre-specified dark-complex target sites, including GAP2 C78, C157, C161; PRK C201, C225; CP12 C60, C69, were highlighted and colored by subunit.

### MED4 and P-HM2 dark complex homology modeling

The MED4 dark complex was modeled with PyMod (v3.0.2)^56^ and Modeller (v10.6)^57^ using the *Thermosynechococcus vestitus* BP-1 dark complex (PDB 6GVE) as template, without symmetry restraints and at MID optimization. Component proteins were extracted from the *P. marinus* subsp. *pastoris* CCMP1986 RefSeq genome (GCA_000011465.1) using our curated annotations^50,51^. P-HM2 CP12 (NCBI YP_004323596.1) was aligned to MED4 CP12 with Clustal Omega (v1.2.4)^58^, adjusted to HMM-matched positions (**Supplementary Methods and Results**).

### Design and assignment of cysteine PTM states in MED4 and P-HM2 dark complexes

Two simulation sets were constructed. A focused redox-state set compared MED4 and P-HM2 CP12 complexes in a fully reduced state and a reference state retaining the CP12 and PRK disulfide bonds modeled from the PDB: 6GVE template^5^. A high-throughput set sampled combinatorial cysteine PTMs across GAP2, PRK, and CP12. Modeled states were *S*-nitrosylation (SNO), *S*-sulfenylation (SOH), *S*-glutathionylation (SSG), reduced thiol (“None”), and, where applicable, disulfide (“SS”). GAP2 carried four modifiable positions (Cys20, Cys78, Cys157, Cys161) across eight chains; PRK carried Cys201 and Cys225 across four chains; CP12 carried two cysteines (MED4 Cys60, Cys69) across four chains, with equivalent positions assigned the same state across copies. Disulfide cysteines were designated CYX and reduced cysteines CYS in the topology. All nonempty subsets of the four GAP2 positions were modified with a single PTM type (15 each for SNO, SOH, SSG), giving 46 GAP2 states with the reduced state. PRK one- and two-site combinations for SNO, SOH, and SSG, plus reduced and disulfide states, gave 11 configurations. CP12 was reduced or disulfide-bonded. Each system combined one GAP2, one PRK, and one CP12 state (46 × 11 × 2 = 1,012 configurations per CP12 background). Oxidative modifications were introduced with PTM-Psi^36^ and disulfides defined explicitly in the topology. The same 1,012 configurations were built for MED4 and P-HM2 CP12 with GAP2, PRK, and all PTM assignments held constant, yielding 2,024 paired systems for direct host-versus-phage comparison.

### Molecular dynamics simulations through the PTM-Psi workflow

Simulations used GROMACS 2025.0^59^ with MPI and GPU acceleration, managed by the Flux scheduler^60,61^ within our PTM-Psi workflow^38^ (v0.1.0; lcf branch for simulations performed on OLCF Frontier and the perlmutter branch for simulations performed on NERSC Perlmutter). Each of the 1,012 configurations followed an identical pipeline. Structures were protonated and parameterized with AMBER99SB^62^, all chains merged, centered in a dodecahedral box (≥1.0 nm clearance), solvated with explicit water (SPC/E), and neutralized to 0.154 M NaCl. Staged energy minimization used position restraints with decreasing force constants (1000, 500, 100, 50, 10, 5, 1 kJ mol⁻¹ nm⁻²), conditionally included in the topology via preprocessor directives, with each step run by steepest descent (integrator = steep, ≤50,000 steps). Systems were then equilibrated at 300 K under NVT for 500 ps with weak protein position restraints (1 kJ mol⁻¹ nm⁻²) and Maxwell–Boltzmann initial velocities (leap-frog, dt = 0.002 ps; velocity-rescaling thermostat, τ = 1.0 ps; nstlist = 10), followed by 500 ps of NPT with the same restraints (isotropic C-rescale barostat, 1 bar, τ = 5.0 ps; velocity-rescaling thermostat, τ = 1.0 ps; nstlist = 20). A final minimization relaxed residual strain, and systems then underwent a second heating/NPT sequence before production. Production MD ran 100 ns (nsteps = 50000000; 2 fs step; cutoff-scheme = Verlet; 1.2 nm electrostatic and van der Waals cutoffs; constraints = h-bonds), with temperature coupling applied separately to the Protein and Non-Protein groups by velocity-rescaling thermostats. Trajectories were water- and ion-stripped, PBC-corrected, and rigid-body aligned to the first frame. The focused analysis used system 0003 (oxidized reference: CP12 Cys60-Cys69 and PRK Cys201-Cys225 as disulfides) and system 0000 (reduced reference: the same cysteines as free thiols, no additional PTMs), giving four conditions (MED4 ox/red, P-HM2 ox/red). Each condition was run as three independent replicates differing only in the random velocity seed (gen-vel = yes, distinct gen-seed) drawn from the same energy-minimized structure. Each replicate was equilibrated independently through the full two-pass heating/NPT protocol before its 100 ns production run, so that thermostat, barostat, and box dimensions equilibrated separately for every replicate. For all statistical comparisons, per-replicate trajectory summaries (per-residue contact frequencies, MM/GBSA energies, BSA/SASA means, contact-dynamics parameters) were treated as independent observations (n = 3 per condition), and individual frames were never pooled across replicates. RMSD, RMSF, radius of gyration, and secondary structure were computed as described^38^.

### CP12 interface analyses

Interface residues were identified from trajectories with MDAnalysis (v2.9.0)^63^ using a heavy-atom contact-frequency approach. For each frame (stride = 1), CP12 (chains C/H/M/P) and partner subunits (GAP2: A/D/E/F/K/I/N/O; PRK: B/G/J/L) were selected without hydrogens, and contacts detected with FastNS at a 5.0 Å cutoff; a residue was in contact when any heavy atom fell within 5.0 Å of a partner heavy atom. Per-residue contact frequency was the fraction of frames with ≥1 contact, stratified as peripheral (≥0.10) or core (≥0.50). Across the four CP12 copies, per-residue frequencies were collapsed by taking the maximum at each aligned position, and copy-robust interfaces required ≥3 copies meeting the peripheral cutoff. Frequencies were aggregated across three replicates as mean ± SEM. To compare MED4 and P-HM2, copy-collapsed frequency vectors were mapped onto the shared CP12 alignment, each column taking the frequency of the corresponding MED4 or P-HM2 residue while system-specific gap positions were treated as missing. For each matched redox state and binding partner, vectors were restricted to columns represented in both systems and compared by Pearson correlation for the CP12-PRK and CP12-GAP2 interfaces; the same procedure applied to within-system oxidized-versus-reduced comparisons to provide a redox baseline for interpreting between-system similarity.

### Contact dynamics analysis

Contact persistence and turnover were quantified using three complementary measures with a 5 Å cutoff. To avoid near-cutoff inflation, a brief disconnect (<300 ps, 3 frames) was not counted as turnover. For each residue pair we computed mean, median, and 90th-percentile dwell time; transitions (0→1, 1→0); turnover rate (events/ns); and two-state k_on and k_off (k_off = 1/⟨τ_on⟩). Pairs were tagged as hydrophobic (both side chains in {A,V,L,I,M,F,W,Y,P}), polar/H-bond (≥1 in {S,T,N,Q,H}), salt-bridge ({D,E}↔{K,R}), cation-π ({K,R}↔{F,W,Y}), or mixed. A per-HMM-position Welch’s t-test on τ was performed with Benjamini-Hochberg FDR correction.

### Buried surface area and solvent accessible surface area

Solvent accessible surface area (SASA)^64^ was computed with gmx sasa (GROMACS 2025.0) from the solvent-stripped, post-processed trajectories, for the whole complex, individual residues, and subunits, including the CP12 pairings (CP12-PRK, CP12-GAP2, CP12-(PRK+GAP2)). Buried surface area between subunits A and B was SASA(A) + SASA(B) − SASA(AB).

### Trajectory principal component analysis and kinetic network analysis

PCA was performed as described^17^ with modifications, analyzing MED4 and P-HM2 ensembles independently and using the reference-state trajectory, which retained the CP12 and PRK disulfide bonds modeled from the 6GVE template, as the reference (labeled “REF”). Eight features were used: mean backbone φ and ψ, radius of gyration, backbone RMSD, mean B-factor from residue fluctuations, and fractional coil, helix, and sheet content. Features were pooled within each ensemble, standardized, and decomposed by full SVD; the smallest set of components explaining ≥90% of variance was retained (six PCs per ensemble; 95.3% MED4, 94.9% P-HM2). Each system was represented by its mean coordinates in PC space and ranked by variance-normalized distance from REF. The top 5% of non-reference systems per ensemble (51 MED4, 51 P-HM2) were retained; 12 PTM configurations appeared in the top 5% of both and were used for matched host-versus-viral comparisons. Mean first-passage time and intra-cluster residence time (τ) were computed as described^17^.

### MM/GBSA binding free energy calculations

Binding free energies were estimated by MM/GBSA using gmx_MMPBSA (v1.6.4)^65^, with CP12 as ligand and GAP2 + PRK + NAD as receptor. To attribute energy to receptor components, additional calculations used CP12 as ligand and each component (PRK or GAP2+NAD) as receptor under identical GB settings and snapshot selection. From each 100 ns solute-only, PBC-corrected trajectory, the first 20 ns were discarded and frames sampled every 400 ps, yielding 200 snapshots. Polar solvation used GB-Neck2 (igb = 8, PBRadii = mbondi3) with the gmx_MMPBSA nonpolar surface-area term, at 300 K and 0.150 M salt with GB-Neck2 default dielectrics. Binding free energy was the Delta (Complex − Receptor − Ligand) term averaged over 200 frames, with PTM effects expressed as ΔG compared to the reference. Per-residue contributions were obtained by MM/GBSA decomposition (idecomp = 2), partitioning 1-4 interactions into electrostatic and van der Waals terms.

## Code availability

Custom scripts and analysis pipelines developed in this study are openly accessible for download at the PNNL NW-BRaVE Zenodo community^66^ under <u>10.5281/zenodo.21536143</u>. The AI agent repository with associated skills is available in the above Zenodo upload.

## Data availability

Primary proteomics raw measurement data files are openly accessible for download at the Mass Spectrometry Interactive Virtual Environment (MassIVE) community repository under the following data accession <u>MSV000102848</u>. Processed redox proteome data, curated CP12 protein family sequence files, and simulation data are openly accessible for download at the PNNL NW-BRaVE Zenodo community under <u>10.5281/zenodo.21536143</u>. Raw molecular dynamics trajectory data have been deposited in EMSL’s Science Central repository and are openly available for download under <u>10.25582/data.2026-08.3856177/3407882</u> (MED4) and <u>10.25582/data.2026-08.3853205/3407881</u> (PHM2). Source data underlying all main and supplementary figures and tables are provided with this paper. All other data supporting the findings of this study are available within the article and its **Supplementary Information** files.

## Acknowledgements

This work was supported by the NW-BRaVE for Biopreparedness project funded by the U.S. Department of Energy (DOE), Office of Science, Office of Biological and Environmental Research, under FWP 81832. Pacific Northwest National Laboratory is a multi-program national laboratory operated by Battelle for the DOE under Contract DE-AC05-76RL01830. Portions of this research were performed at the Environmental Molecular Sciences Laboratory, a DOE Office of Science User Facility sponsored by the Biological and Environmental Research program, under Proposal 61054 (Award DOI: 10.46936/staf.proj.2023.61054/60012367) and the Tahoma HPC facility. NW-BRaVE additionally leveraged computational resources from the Advanced Computing Ecosystem IRI testbed at Oak Ridge National Laboratory’s Leadership Computing, which is a DOE Office of Science User Facility supported under Contract DE-AC05-00OR22725 and the National Energy Scientific Computing Center (NERSC), a Department of Energy User Facility using NERSC award ALCC-ERCAP0034213 and under Contract No. DE-AC02-05CH11231 using NERSC award BER-ERCAP0038634. The research described herein was funded by the Generative AI for Science, Energy, and Security Science & Technology Investment under the Laboratory Directed Research and Development Program at Pacific Northwest National Laboratory (PNNL). This work was also supported by the Center for AI and Center for Cloud Computing at PNNL.

## Author contributions

M.S.C., K.I., and W.J.Q. designed the research. S.W., R.W., M.N. and A.A. performed the computational research. S.W., R.W., M.N, A.A., K.I., and M.S.C. contributed to the analysis of the simulation data. D.M.R., S.S., R.N., H.K., A.G., and P.R. contributed to the computational workflow optimization. P.B. and X.L. performed experimental work to generate omics data. J.R., S.F., P.B., R.W., N.B. contributed the analysis of the multimodal data for the computational research. L.N.A., R.W., M.N. and A.G. packaged the multimodal data and analysis scripts for release. S.W., M.N., R.W., A.A., M.S.C. wrote the first draft of the paper. All authors contributed to writing the paper.

## Supplementary Figures

**Supplementary Figure 1. CP12 structural-functional features among metagenome-derived and reference sequences vary across taxonomic lineages.** The distribution of three CP12 structural-functional features was compared across full-length metagenome-derived CP12 sequences, partial metagenome-derived CP12 sequences, and reference CP12 sequences. Source-dependent distributions for the number of cysteine residues (**a**), mean cysteine spacing (**b**), and acidic residue fraction (**c**), with violins showing the overall distribution and boxplots showing the median and interquartile range. Group sizes are indicated below each source category. Significance was assessed using pairwise Wilcoxon rank-sum tests, with adjusted significant comparisons shown above the plots (< 0.05, *; < 0.0001, ****). **d** lineage-resolved density distributions for the same three features, stratified by source group (full sequences for metagenome in green, partial sequences from metagenome in orange, and reference sequences extracted from public databases in blue) and faceted by major taxonomic lineage. The number of sequences represented in each lineage is shown in the upper-right corner.

**Supplementary Figure 2. Null model analysis of chemistry-level constraint across CP12 interface classes.** Observed values (colored points) are compared against null distributions (gray points and boxplots) generated by randomly reassigning amino acids to chemical classes while preserving amino acid frequencies. Boxplots show the median, interquartile range, and whiskers of the permutation-derived null distributions for each interface class. The top panel shows chemical class dominance represented by the maximum summed fraction of residues belonging to any one chemical class, noted as pmax_Chem, and the bottom panel shows the chemistry-minus-identity gap (Δ = pmax_Chem-pmax_AA; pmax_AA is the maximum amino acid fraction at that position). Points above the null distribution indicate enrichment beyond what is expected from random category assignment. P-values indicate the fraction of null permutations exceeding the observed median for each interface class.

**Supplementary Figure 3. Physicochemical property changes in viral CP12 relative to host CP12.** Physicochemical property changes at each amino acid substitution between native MED4 and viral HM2 CP12, plotted along the Pfam PF02672 HMM coordinate system. Three properties are shown: charge change (top), hydrophobicity change (Kyte-Doolittle scale, middle), and volume change (van der Waals volume in Å³, bottom; Richards-1974 average residue volume). Red bars indicate positive changes (increased charge, hydrophobicity, or volume in HM2 relative to MED4); blue bars indicate negative changes. The aligned MED4 and HM2 sequences are displayed below the property plots, with substituted positions highlighted in yellow. Charge was computed as the difference in net charge at pH 7.0 (K/R = +1, H = +0.5, D/E = −1, all others = 0). Hydrophobicity was calculated from the Kyte-Doolittle scale. Volume changes were computed from Richards (1974) average residue volumes. Substitutions were identified from the PF02672 profile-HMM alignment of MED4 (74 residues) and HM2 (70 residues) CP12 sequences (HMMER hmmalign v3.3.2; 70 HMM match-state columns; 34 conserved, 31 substituted, 5 gap positions; 52.3% identity over the 65 aligned positions). The largest charge changes are at K12E (−2.0), E16K (+2.0), and D13H (+1.5); the largest hydrophobicity changes at V29R (−8.7), Q14I (+8.0), I17D (−8.0), K21L (+7.7), and K28A (+5.7); the largest volume changes at K28A (−80.0 Å³), G41V (+79.9), A20I (+78.1), A62M (+74.3), and A18H (+64.6). Twenty of 31 substitutions (65%) lie at PRK-interface positions, one at the GAP2 interface, and ten at non-interface positions. MED4 N-terminal residues 1–5 (MKKKS) and the C-terminal Asp lie outside the HMM model and are not shown on the alignment-position axis.

**Supplementary Figure 4. Performance matrix of the viral versus bacterial CP12 classifier**. **a** Receiver operating characteristic (ROC) curve based on HMM position-resolved chemical class features. The dashed diagonal line indicates random performance, and the shaded region indicates the bootstrap 95% confidence interval. **b** Precision-recall (PR) curve showing performance under class imbalance. The dashed line indicates the random baseline and the shaded region indicates the bootstrap 95% confidence interval. **c** Confusion matrix from out-of-fold predictions at a probability threshold of 0.5, showing high viral recall and bacterial specificity.

**Supplementary Figure 5. Homology modeling and assessment. a** Ramachandran plots of the MED4 and P-HM2 dark complex models. Before (top) and after (bottom) energy minimization. φ/ψ angles for all complex residues, colored as favored (blue), allowed (orange), or outlier (red); dotted boxes denote favored regions. ≥95 % of residues lie in favored regions in all four structures (≤2.7 % outliers). **b** Backbone RMSD (Å) for reference systems of MED4 (blue) and P-HM2 (orange) over the 100ns trajectory. The shaded area is 1 standard deviation (SD) over three replicates. **c** Relative binding free energies associated with disulfide bond formation for cysteine pairs in HM2 and MED4. Orange bars represent the Cys56/Cys65 disulfide pair in subunits C, H, M, and P of HM2, while blue bars correspond to the structurally homologous Cys60/Cys69 pair in MED4. Error bars indicate estimated uncertainties from the free energy calculations.

**Supplementary Figure 6. CP12 RMSF by interface region.** Mean CP12 C-alpha RMSF for PRK-binding and GAP2-binding regions in MED4 and P-HM2 CP12 under (a) oxidized and (b) reduced conditions. Bars show mean RMSF with SEM. Host comparisons within each region and redox state were tested with two-sided Welch’s t-tests; asterisk denotes p < 0.05, and ns denotes not significant.

**Supplementary Figure 7. P-HM2 CP12 preserves total Dark Complex stability through position-localized VdW contacts at sequence-divergent residues. a** Total MM/GBSA binding energy for CP12 binding to GAP2/NAD/PRK. **b** Total MM/GBSA binding energy for CP12 binding to PRK. **c** Total MM/GBSA binding energy for CP12 binding to GAP2/NAD. d Per-position MM/GBSA VdW energy at the CP12–PRK interface for MED4 (blue) and HM2 (orange) CP12, oxidized (left) and reduced (right). Each point is one CP12 alignment column, averaged across four chain copies and three replicates. Positions are split by category: substituted (n = 31, MED4 ≠ HM2) and conserved (n = 34, identical residue). Lower brackets: within-category Welch’s t-tests (MED4 vs HM2). Upper bracket: interaction test (Welch’s t on per-position HM2 - MED4 differences, substituted vs conserved). **e** ΔVdW = HM2 - MED4 (kcal/mol per chain) at each CP12 alignment column for the CP12-PRK interface, oxidized (top) and reduced (bottom). Each marker is one position, averaged across four CP12 chain copies and three replicates. Substituted positions (MED4 ≠ HM2 residue) are diamonds (orange = HM2 advantage, blue = MED4 advantage); conserved positions are grey circles. In the reduced panel, the five substituted positions with the largest |ΔVdW| are labeled.

**Supplementary Figure 8.** CP12 contact dynamics reveal partner-specific remodeling across system and redox comparisons. CP12 contact-dynamics differences across PRK and GAP2 interfaces, comparing P-HM2 against MED4 under oxidized and reduced conditions and comparing reduced against oxidized states within each CP12 system. CP12 HMM-aligned positions are labeled as MED4/P-HM2 residue pairs and colored by residue-chemistry shift. The heatmaps summarize contact behavior from 1001 trajectory frames sampled every 100 ps for each replicate, using 5 Å heavy-atom CP12-partner contacts to track how many partner residues each CP12 position contacted (No. contacts), how close those contacts were (Contact dist.), how often they occurred (contact frequency), how long they persisted (log10 dwell time), and how frequently they turned over (contact turnover per ns). For system comparisons, values represent P-HM2 minus MED4. For redox comparisons, values represent reduced minus oxidized within MED4 or P-HM2. Values were calculated within matched replicates before being summarized across replicate pairs and scaled within each contact-dynamics metric. Red indicates higher values in P-HM2 or reduced simulations, blue indicates higher values in MED4 or oxidized simulations, and near-white indicates little difference. Gray indicates that the metric was not estimable, typically because no contact was detected in one or both compared conditions. Symbols mark one-sided data availability with upward triangles indicating contacts present only in P-HM2 or reduced simulations, downward triangles indicating contacts present only in MED4 or oxidized simulations, and gray circles indicating non-estimable values. Asterisks indicate statistical support for non-zero paired differences in the heatmap metrics. For each CP12 HMM position, binding partner, comparison, and metric, replicate-level deltas were tested against zero using a one-sample t-test across three matched replicate simulations. Significance levels are shown as p < 0.05, p < 0.01, and p < 0.001.

**Supplementary Figure 9. Residue-chemistry dependence of P-HM2 versus MED4 contact-dynamics shifts.** Distributions show the differences in P-HM2 minus MED4 Dark Complex contact-dynamics parameters, grouped by residue-chemistry class and separated by redox state and binding partner. Each boxplot summarizes HMM-position-level shift values for the indicated residue-chemistry class. Points represent individual HMM positions, boxes show the interquartile range, center lines show medians, and whiskers show the plotted data range. Positive values indicate higher values in P-HM2 than MED4, whereas negative values indicate lower values in P-HM2 than MED4. Sample sizes for each residue-chemistry class are shown below the boxes. Residue-chemistry classes include conserved positions, within-class substitutions, Gly/Pro gain or loss, hydrophobic-polar substitutions, hydrophobic/polar-charged substitutions, and charge-flip substitutions. Kruskal-Wallis tests evaluate whether P-HM2 minus MED4 shift distributions differ among residue-chemistry classes within each panel. Asterisks indicate the pairwise post hoc differences with significant residue-class effects highlighted.

**Supplementary Figure 10.** BSA and SASA dynamics reveal selective remodeling of the GAP2-facing interface. Time-resolved buried surface area (BSA, a) and solvent-accessible surface area (SASA, b) profiles compare MED4 (blue line) and P-HM2 (orange line) Dark Complex simulations under oxidized and reduced conditions. Thin lines represent individual replicate trajectories, and thick lines represent the system-level mean. Columns show oxidized and reduced simulations; rows show CP12 interaction with all partners, PRK, or GAP2. P values from paired t-test comparing MED4 and P-HM2 replicate-level means within each redox state; asterisks indicate p < 0.05.

**Supplementary Figure 11. Conformational displacement scales with PTM load. a** Variance-normalized PCA distance to the unmodified reference as a function of total PTM count. MED4 (blue) versus P-HM2 (orange). Each box summarizes all systems carrying the indicated number of PTMs (median, interquartile range; whiskers, 1.5 IQR). MED4 Spearman ρ = 0.688, p = 2.2 × 10⁻¹⁴²; P-HM2 ρ = 0.696, p = 1.3 × 10⁻¹⁴⁷; correlations computed on per-system values, not the binned boxes. **b** Subunit-resolved correlation between PTM count and conformational divergence. Spearman ρ between the number of PTMs on each subunit (GAP2, PRK, CP12) and variance-normalized PCA distance to the unmodified reference, for MED4 (blue) and P-HM2 (orange); points are ρ, horizontal bars the 95% confidence interval. Asterisks denote the correlation p-value (**** p < 10⁻⁴, * p < 0.05).

**Supplementary Figure 12. PTM enrichment among the most displaced MED4 and P-HM2 systems. a** Enrichment of each PTM chemistry among the most displaced systems. Log₂ fold enrichment (top-5% versus background prevalence) of *S*-glutathionylation (SSG), sulfenylation (SOH), and *S*-nitrosylation (SNO) in MED4 (blue) and P-HM2 (orange); positive values indicate over-representation in the top 5%. Asterisks denote significance (**** q < 10⁻⁴, * q < 0.05; BH-adjusted Mann–Whitney on per-system PTM count). b Site- and chemistry-resolved PTM enrichment in the top 5% most displaced systems. Each row is a single PTM feature (chemistry × residue); x-position gives its prevalence among the top-5% systems (PTM ratio), dot color the enrichment significance (−log₁₀ BH-adjusted Fisher q), and dot size the number of top-5% systems carrying the feature, for MED4 (left) and P-HM2 (right). The strongest feature in both ensembles is SSG of GAP2 Cys20, and the enriched signal localizes to the GAP2 N-terminal cysteine cluster (Cys20, 78, 157, 161).

**Supplementary Figure 13. Matched-pair conformational kinetics of the 12 shared top-5% systems under MED4 versus P-HM2 CP12. a** MFPT and **b** intra-cluster residence time τ as log₂ ratios to the ensemble reference; paired MED4 (blue) and P-HM2 (orange) values joined by a line, dashed line = no change.

**Supplementary Figure 14. Viral CP12 buffers the PTM-induced loss of CP12 binding affinity. a** MM/GBSA binding free energy of CP12 (4 chains) to the GAP2+PRK+NAD receptor, as change vs each ensemble’s own reference (ΔΔG vs REF); dotted line = reference. MED4 blue, P-HM2 orange. **A** Distribution over the 51 top systems (top 5%) per ensemble (violin + points, bar = median); Mann-Whitney p = 5×10⁻⁸. **b** The 12 shared systems (Overlapping 12; identical GAP2+PRK PTMs) as matched host-viral pairs (bar = mean); paired Wilcoxon p = 0.001.

**Supplementary Movie 1.** An all-atom **t**rajectory movie comparing MED4 (left) and P-HM2 (right) reference oxidized system 0003. Specific subunits highlighted are GAP2 chain O (gray), PRK chain L (purple), and CP12 chain M (cyan). MED4 exhibits a compacted overall structure, while the P-HM2 viral version exhibits an expanded structure with greater engagement at the PRK interface.

**Supplementary Movie 2.** Spatial principal component analysis is computed via normal mode over each reference oxidized trajectory (0003) and visualized for MED4 (left), and P-HM2 (right). The modes with greatest displacement (MED4: modes 2 and 8, P-HM2: modes 4 and 10) between GAP2 (gray) and PRK (purple) are used. A greater flexibility can be localized to the CP12 chains in the viral complex, and viral CP12 retains solvent exposure to a greater degree, as do the GAP2 subunits of the viral complex.

## References

1. Gurrieri, L., Fermani, S., Zaffagnini, M., Sparla, F. & Trost, P. Calvin–Benson cycle regulation is getting complex. Trends Plant Sci. 26, 898–912 (2021).

2. Hohmann-Marriott, M. F. & Blankenship, R. E. Evolution of photosynthesis. Annu. Rev.Plant Biol. 62, 515–548 (2011).

3. Marri, L. et al. Spontaneous assembly of photosynthetic supramolecular complexes as mediated by the intrinsically unstructured protein CP12. J. Biol. Chem. 283, 1831–1838 (2008).

4. Graciet, E., Lebreton, S. & Gontero, B. Emergence of new regulatory mechanisms in the Benson–Calvin pathway via protein–protein interactions: a glyceraldehyde-3-phosphate dehydrogenase/CP12/phosphoribulokinase complex. J. Exp. Bot. 55, 1245–1254 (2004).

5. McFarlane, C. et al. Structural basis of light-induced redox regulation in the Calvin– Benson cycle in cyanobacteria. Proc Natl Acad Sci 116, 20984–20990 (2019).

6. Flombaum, P. et al. Present and future global distributions of the marine Cyanobacteria Prochlorococcus and Synechococcus. Proc. Natl. Acad. Sci. 110, 9824–9829 (2013).

7. Biller, S. J., Berube, P. M., Lindell, D. & Chisholm, S. W. Prochlorococcus: the structure and function of collective diversity. Nat. Rev. Microbiol. 13, 13–27 (2015).

8. Suttle, C. A. Marine viruses — major players in the global ecosystem. Nat. Rev. Microbiol. 5, 801–812 (2007).

9. Johnson, C. G. et al. Multi-omics reveals temporal scales of carbon metabolism in synechococcus elongatus PCC 7942 under light disturbance. PRX Life 3, 033017 (2025).

10. Rozum, J. C. et al. Synergy and antagonism in a genome-scale model of metabolic hijacking by bacteriophages. Sci. Adv. 12, eaeb7646 (2026).

11. Thompson, L. R., et al. Phage auxiliary metabolic genes and the redirection of cyanobacterial host carbon metabolism. Proc. Natl. Acad. Sci. 108, E757–E764 (2011).

12. Cai, L., Li, H., Deng, J., Zhou, R. & Zeng, Q. Biological interactions with Prochlorococcus: implications for the marine carbon cycle. Trends Microbiol. 32, 280–291 (2024).

13. Thompson, L., Zeng, Q. & Chisholm, S. Gene Expression Patterns during Light and Dark Infection of Prochlorococcus by Cyanophage. PLoS ONE 11, (2016).

14. Jumper, J., Evans, R. & Pritzel, A. Highly accurate protein structure prediction with AlphaFold. Nature 596, (2021).

15. Hollingsworth, S. A. & Dror, R. O. Molecular Dynamics Simulation for All. Neuron 99, 1129–1143 (2018).

16. Li, H. & Ma, A. Enhanced sampling of protein conformational changes via true reaction coordinates from energy relaxation. Nat. Commun. 16, 786 (2025).

17. Samantray, S. et al. PTM-Psi on the Cloud: A Cloud-Compatible Workflow for Scalable, High-Throughput Simulation of Post-Translational Modifications in Protein Complexes. J. Chem. Inf. Model. 65, 11473–11485 (2025).

18. Childers, M. C. & Daggett, V. Insights from molecular dynamics simulations for computational protein design. Mol. Syst. Des. Eng. 2, 9–33 (2017).

19. Guo, J. et al. Proteome-wide Light/Dark Modulation of Thiol Oxidation in Cyanobacteria Revealed by Quantitative Site-specific Redox Proteomics. Mol. Cell. Proteomics 13, 3270–3285 (2014).

20. Mallén-Ponce, M. J., Florencio, F. J. & Huertas, M. J. Thioredoxin A regulates protein synthesis to maintain carbon and nitrogen partitioning in cyanobacteria. Plant Physiol. 195, 2921–2936 (2024).

21. Gurrieri, L., Sparla, F., Zaffagnini, M. & Trost, P. Dark complexes of the Calvin-Benson cycle in a physiological perspective. Semin. Cell Dev. Biol. 155, 48–58 (2024).

22. Howard, T. P., Metodiev, M., Lloyd, J. C. & Raines, C. A. Thioredoxin-mediated reversible dissociation of a stromal multiprotein complex in response to changes in light availability. Proc. Natl. Acad. Sci. U. S. A. 105, 4056–4061 (2008).

23. Boise, N. R. et al. Sampling microbial dynamics in the Salish Sea estuary: evaluating methods to capture cyanobacteria and cyanophage. Front. Mar. Sci. 13, 1769457 (2026).

24. Kim, H. et al. Thiol post-translational modifications modulate allosteric regulation of the OpcA–G6PDH complex through conformational gate control. Protein Sci. 35, e70561 (2026).

25. Mann, N., Cook, A., Millard, A., Bailey, S. & Clokie, M. Bacterial photosynthesis genes in a virus. Nature 424, 741 (2003).

26. López-Calcagno, P. E., Howard, T. P. & Raines, C. A. The CP12 protein family: a thioredoxin-mediated metabolic switch? Front. Plant Sci. 5, (2014).

27. Gurrieri, L., Sparla, F., Zaffagnini, M. & Trost, P. Dark complexes of the Calvin-Benson cycle in a physiological perspective.

28. Tripathi, S., Waxham, M. N., Cheung, M. S. & Liu, Y. Lessons in Protein Design from Combined Evolution and Conformational Dynamics. Sci. Rep. 5, 14259 (2015).

29. Frank, J. A. et al. Structure and function of a cyanophage-encoded peptide deformylase. ISME J. 7, 1150–1160 (2013).

30. Forrey, C., Douglas, J. F. & Gilson, M. K. The fundamental role of flexibility on the strength of molecular binding. Soft Matter 8, 6385–6392 (2012).

31. Zhao, F. et al. Biochemical and structural characterization of the cyanophage-encoded phosphate-binding protein: implications for enhanced phosphate uptake of infected cyanobacteria. Environ. Microbiol. 24, 3037–3050 (2022).

32. Lucius, S. et al. CP12 fine-tunes the Calvin-Benson cycle and carbohydrate metabolism in cyanobacteria. Front. Plant Sci. 13, (2022).

33. Zinser, E. Choreography of the Transcriptome, Photophysiology, and Cell Cycle of a Minimal Photoautotroph, Prochlorococcus. 4, e5135 (2009).

34. Waldbauer, J. R., Rodrigue, S., Coleman, M. L. & Chisholm, S. W. Transcriptome and Proteome Dynamics of a Light-Dark Synchronized Bacterial Cell Cycle. PLoS One 7, (2012).

35. Demory, D. Linking Light-Dependent Life History Traits with Population Dynamics for Prochlorococcus and Cyanophage. mSystems 5, (2020).

36. Zaffagnini, M. et al. The thioredoxin-independent isoform of chloroplastic glyceraldehyde-3-phosphate dehydrogenase is selectively regulated by glutathionylation. FEBS J. 274, 212–226 (2007).

37. Chardonnet, S. et al. First Proteomic Study of S-Glutathionylation in Cyanobacteria. J. Proteome Res. 14, 59–71 (2015).

38. Mejia-Rodriguez, D. et al. PTM-Psi: A python package to facilitate the computational investigation of post-translational modification on protein structures and their impacts on dynamics and functions. Protein Sci. 32, (2023).

39. González, M. M., Abriata, L. A., Tomatis, P. E. & Vila, A. J. Optimization of Conformational Dynamics in an Epistatic Evolutionary Trajectory. Mol. Biol. Evol. 33, 1768–1776 (2016).

40. Paulsen, C. E. & Carroll, K. S. Cysteine-mediated redox signaling: chemistry, biology, and tools for discovery. Chem. Rev. 113, 4633 (2013).

41. Swint-Kruse, L. & Fenton, A. W. Rheostats, toggles, and neutrals, Oh my! A new framework for understanding how amino acid changes modulate protein function Author links open overlay panel. J. Biol. Chem. 300, (2024).

42. Wright, P. E. & Dyson, H. J. Intrinsically Disordered Proteins in Cellular Signaling and Regulation. Nat. Rev. Mol. Cell Biol. 16, 18–29 (2015).

43. Bah, A. & Forman-Kay, J. D. Modulation of Intrinsically Disordered Protein Function by Post-translational Modifications. J. Biol. Chem. 291, 6696–6705 (2016).

44. Bohutskyi, P. et al. Systematic scale-up and enhanced purification of marine cyanophage P-SSP7. Front. Microbiol. 17, 1776133 (2026).

45. Gluth, A. et al. Integrative Multi-PTM Proteomics Reveals Dynamic Global, Redox, Phosphorylation, and Acetylation Regulation in Cytokine-Treated Pancreatic Beta Cells. Mol. Cell. Proteomics MCP 23, 100881 (2024).

46. Day, N. J. et al. A deep redox proteome profiling workflow and its application to skeletal muscle of a Duchenne Muscular Dystrophy model. Free Radic. Biol. Med. 193, 373–384 (2022).

47. Guo, J. et al. Resin-assisted enrichment of thiols as a general strategy for proteomic profiling of cysteine-based reversible modifications. Nat. Protoc. 9, 64–75 (2014).

48. Gaffrey, M. J., Day, N. J., Li, X. & Qian, W.-J. Resin-Assisted Capture Coupled with Isobaric Tandem Mass Tag Labeling for Multiplexed Quantification of Protein Thiol Oxidation. J. Vis. Exp. JoVE 10.3791/62671 (2021) doi:10.3791/62671.

49. Kim, S. & Pevzner, P. A. MS-GF+ makes progress towards a universal database search tool for proteomics. Nat. Commun. 5, 5277 (2014).

50. Cheung, M. S., Feng, S. & Qian, W.-J. MassIVE MSV000100371 - Time-Course Sampled Axenic Prochlorococcus marinus MED4 Proteomics: Prochlorococcus marinus MED4 uninfected axenic cell cultures for LC-MS proteome analysis of time sampled circadian experiments. Peptides were extracted using the MPLEx protocol (PubMed: 27822525) from cell pellet samples and prepared for DIA for LC-MS. Data was searched with DIA-NN using PNNL’s DMS processing pipeline. Normalized protein counts (log2) were generated using intensity-based absolute quantification (iBAQ) for downstream flux balance analysis. Processed results files have been packaged under the NW-BRaVE project dataset DOI: 10.5281/zenodo.18022273. MassIVE 10.25345/C5KH0FC46 (2026).

51. Feng, S., et al. Circadian Time-Course Sampled Axenic Prochlorococcus marinus MED4 Multi-Omics. Zenodo 10.5281/ZENODO.18022272 (2026).

52. Petyuk, V. PlexedPiper. Zenodo 10.5281/ZENODO.14708767 (2025).

53. Rozum, J. C. et al. ProteoMeter: A pipeline for integrating multi-PTM and limited proteolysis data to reveal modification-structure coupling at the residue level. Preprint at 10.1101/2025.07.14.664747 (2025).

54. Agent Skills contributors. Agent Skills.

55. Code, V. S. Visual studio code. Recuperado El Oct. De (2019).

56. Janson, G. & Paiardini, A. PyMod 3: a complete suite for structural bioinformatics in PyMOL. Bioinformatics 37, 1471–1472 (2021).

57. Sali, A. & Blundell, T. L. Comparative protein modelling by satisfaction of spatial restraints. J. Mol. Biol. 234, 779–815 (1993).

58. Sievers, F. et al. Fast, scalable generation of high-quality protein multiple sequence alignments using Clustal Omega. Mol. Syst. Biol. 7, 539 (2011).

59. Van Der Spoel, D. et al. GROMACS: fast, flexible, and free. J. Comput. Chem. 26, 1701–1718 (2005).

60. Ahn, D. et al. Flux: A next-generation resource management framework for large HPC centers. 2014 43rd Int. Conf. Parallel Process. Workshop (2014).

61. Ahn, D. et al. Flux: Overcoming scheduling challenges for exascale workflows. Future Gener. Comput. Syst. 110, 202–213 (2020).

62. Hornak, V. et al. Comparison of multiple Amber force fields and development of improved protein backbone parameters. Proteins 65, 712–725 (2006).

63. Michaud-Agrawal, N., Denning, E. J., Woolf, T. B. & Beckstein, O. MDAnalysis: A Toolkit for the Analysis of Molecular Dynamics Simulations. J. Comput. Chem. 32, 2319–2327 (2011).

64. Eisenhaber, F., Lijnzaad, P., Argos, P., Sander, C. & Scharf, M. The double cubic lattice method: Efficient approaches to numerical integration of surface area and volume and to dot surface contouring of molecular assemblies. J. Comput. Chem. 16, 273–284 (1995).

65. Valdés-Tresanco, M. S., Valdés-Tresanco, M. E., Valiente, P. A. & Moreno, E. gmx_MMPBSA: A New Tool to Perform End-State Free Energy Calculations with GROMACS. J. Chem. Theory Comput. 17, 6281–6291 (2021).

66. Nelson, Malio et al. Analysis Scripts for Cyanophage CP12 Rewires Host Carbon Regulation through Interface Remodeling and Redox Buffering. Zenodo 10.5281/zenodo.21536143.

